# Engineering single-AAV CRISPR-Cas13 RNA base editors for treatment of inherited retinal diseases

**DOI:** 10.1101/2025.08.06.668808

**Authors:** Satheesh Kumar, Heng-Ai Chang, Deborah Aubin, Yi-Wen Hsiao, Alicia Brunet, Jun Yang, Liuhui Huang, Chi D Luu, Alex W Hewitt, Fan Li, Lewis E Fry, Livia S Carvalho, Anai Gonzalez-Cordero, Guei-Sheung Liu

**Author notes:** Correspondence and requests for materials should be addressed to Dr Guei-Sheung Liu. Centre for Eye Research Australia. Address: Level 10, 200 Victoria Parade, East Melbourne, VIC 3002, Australia.

## Abstract

Site-directed RNA editing, especially RNA base editing, allows for specific manipulation of RNA sequences, making it a useful approach for the correction of pathogenic mutations. Correction of RNA transcripts allows therapeutic gene editing in a safe and reversible manner and avoids permanent alterations in the genome. RNA-targeting CRISPR-Cas nucleases (e.g., CRISPR-Cas13) enable delivery within a single adeno-associated virus (AAV) vector for RNA base editing, making the approach clinically feasible. Here, we used the inactive CRISPR-Cas13bt3 (also known as Cas13X.1) fused to the ADAR2 deaminase domain (ADAR2DD) for targeted correction of inherited retinal disease (IRD) mutations. First, we show *in vitro* that dCas13bt3-ADAR2DD can efficiently correct a pathogenic nonsense mutation (c.130C>T [p.R44X]) found in the mouse *Rpe65* gene and recover protein expression in retinal pigment epithelium cells (RPEs). Across clinically reported *RPE65* mutations, we observed editing efficiencies ranging from 0% to 60%. In the *Rpe65*-deficient mouse model of retinal degeneration (rd12), we observed that RNA base editing can recover *Rpe65* expression in RPEs and rescue retinal function with no observable adverse effects. We further employed our RNA base editor against the large *USH2A* gene to assess the promise of RNA base editing for addressing untreatable IRDs caused by genes too large for AAV gene delivery. Against the human *USH2A in vitro*, we observed up to 60% on-target efficiency. We further found that gRNA mismatches, domain-inlaid ADAR2DD design and nucleocytoplasmic shuttling of the RNA base editor optimised on-target and bystander editing for a highly precise base editor. Against the mouse *Ush2a in vitro*, we similarly observed up to 60% on-target editing in mammalian cells, while in the *Ush2a*^W3947X^ mice, we observed ∼12% on-target editing, with no impact on retinal structure or function, or transcriptome-wide editing. Overall, our findings demonstrate dCas13bt3-ADAR2DD as a potent tool for gene therapy against IRDs, addressing a significant unmet clinical need in ophthalmology.

## INTRODUCTION

Inherited retinal diseases (IRD) collectively are a leading cause of blindness in working-age adults globally^1^. Affecting 1 in 2000 individuals, they can be caused by mutations that occur in over 300 genes^2^. To date, only one form of IRD, *RPE65*-associated Leber Congenital Amaurosis (LCA), has an approved therapy. Branded Luxturna (voretigene-neparvovec), it became the first *in vivo* gene therapy to be approved by the United States Food and Drug Administration (US FDA) in 2017. The development of similar therapies for other forms of IRDs, however, has been challenging due to the size limitation of adeno-associated viral (AAV) vector. An analysis of IRD-causing genes as reported in the Leiden Open Variation Database (LOVD), revealed that over to 50% of pathogenic variants were editable by currently established base editors^3^.

Clustered regularly interspaced short palindromic repeats (CRISPR) and CRISPR-associated (Cas) technology has accelerated gene therapy by enabling treatments for familial hypercholesteremia, Duchenne muscular dystrophy, transthyretin amyloidosis, Hutchinson-Gilford progeria and Leber congenital amaurosis^4, 5, 6, 7^, using a combination of approaches such as targeted gene knockdown, activation, repression, base editing and prime editing^8, 9, 10^. Recent advances in RNA base editing with CRISPR-Cas13 nucleases have also been shown to provide therapeutic benefits without changing the genome, and compact versions enable delivery with a single AAV vector^11, 12^. While some Cas13 enzymes (i.e., Cas13a) may require protospacer flanking sites (PFS), Cas13b and the related Cas13bt enzymes are not restricted by the PFS^13, 14^, promising a widely applicable tool. AAV-delivered RNA base editors can therefore potentially target a wide range of mutations, across many IRD genes.

To this end, we developed a Cas13-mediated RNA base editor using the compact Cas13bt3 nuclease^15^. This proved more efficient than truncated versions of PspCas13b (used in previously reported REPAIR systems^16^), and the compact and human protein-based CRISPR-inspired RNA targeting system (CIRTS)^17^. It also proved effective at reversing the mutation across a variety of clinically occurring *RPE65* mutations. We further demonstrate ∼2-3% editing of *Rpe65* mRNA in the retinal degeneration 12 (rd12) mouse model, leading to the rescue of cone photoreceptor function. We then targeted the *Ush2a*^W3947X^ variant orthologous to the most common pathogenic G>A mutation in *USH2A*, c.11864G>A (W3955X), where over 50% correction in mRNA was observed *in vitro*. Intravitreal delivery of AAV2.7m8 viruses into *Ush2a*^W3947X^ mice resulted in a ∼10-20% correction in mRNA, and the recovery of Usherin in photoreceptors. Against the human *USH2A*^W3955X^, we demonstrate high on-target editing with significant bystander editing. We describe gRNA and Cas13-ADAR fusion protein engineering methods as well as the incorporation of nucleocytoplasmic shuttling to optimise editing efficiency and specificity. Overall, our results demonstrate that RNA base editing is a potent and versatile tool with wide applicability to several IRD mutations. The *in vivo* efficacy also shows promise for a new gene therapy against IRDs, particularly those caused by large genes that cannot be treated by gene replacement.

## RESULTS

### AAV-delivered RNA base editors can address a significant unmet clinical need

To determine the applicability of AAV-deliverable gene editing tools for IRDs, we analysed IRD genes and variants reported in the LOVD database. Approximately 50% of genes implicated in IRDs exceeded the AAV carrying capacity (approximately 3.8 kb, accounting for promoter and terminator requirements) and cannot be delivered by gene replacement therapies like Luxturna (**Figure S1a**). Point mutations account for over 70% of variants in these genes (**Figure S1b**), making base editing a promising approach to address a broad spectrum of IRDs. Specific to RNA base editing, currently available tools can edit approximately 48% of point mutation variants, and over 30% of point mutations can be precisely reversed using RNA base editing tools to induce A>G or C>U changes (**Figure S1c-d**). An AAV-mediated RNA base editor would therefore offer a safe and versatile tool for addressing a wide range of IRD mutations.

### dCas13bt3-ADAR2DD efficiently corrects the Rpe65^R44X^ mutation and recovers protein expression

To develop an all-in-one AAV RNA base editor, we truncated 150 nucleotides (nt) from inactive Cas13bt3 (dCas13bt3) at the C-terminal as previously described^15^, and fused the ADAR2 deaminase domain (ADAR2DD). This was flanked with a U6-driven 50-nt gRNA with a centrally placed (at the 25^th^ position) A:C mismatch (**Figure 1a-b**) targeting the nonsense mutation (G>A) of the mCherry reporter gene (**Figure S2**). We benchmarked the truncated dCas13bt3-ADAR2DD base editor to the HEPN-domain truncated dPspCas13b-ADAR2DD base editor and the CRISPR-inspired RNA-targeting system (CIRTS)^17^, a compact human protein-based RNA base editor (**Figure S2b-d**). Our results demonstrated that the dCas13bt3-ADAR2DD base editor was significantly more efficient in restoring mCherry expression compared to the other two systems. (**Figure S2b**).

**Figure 1.**
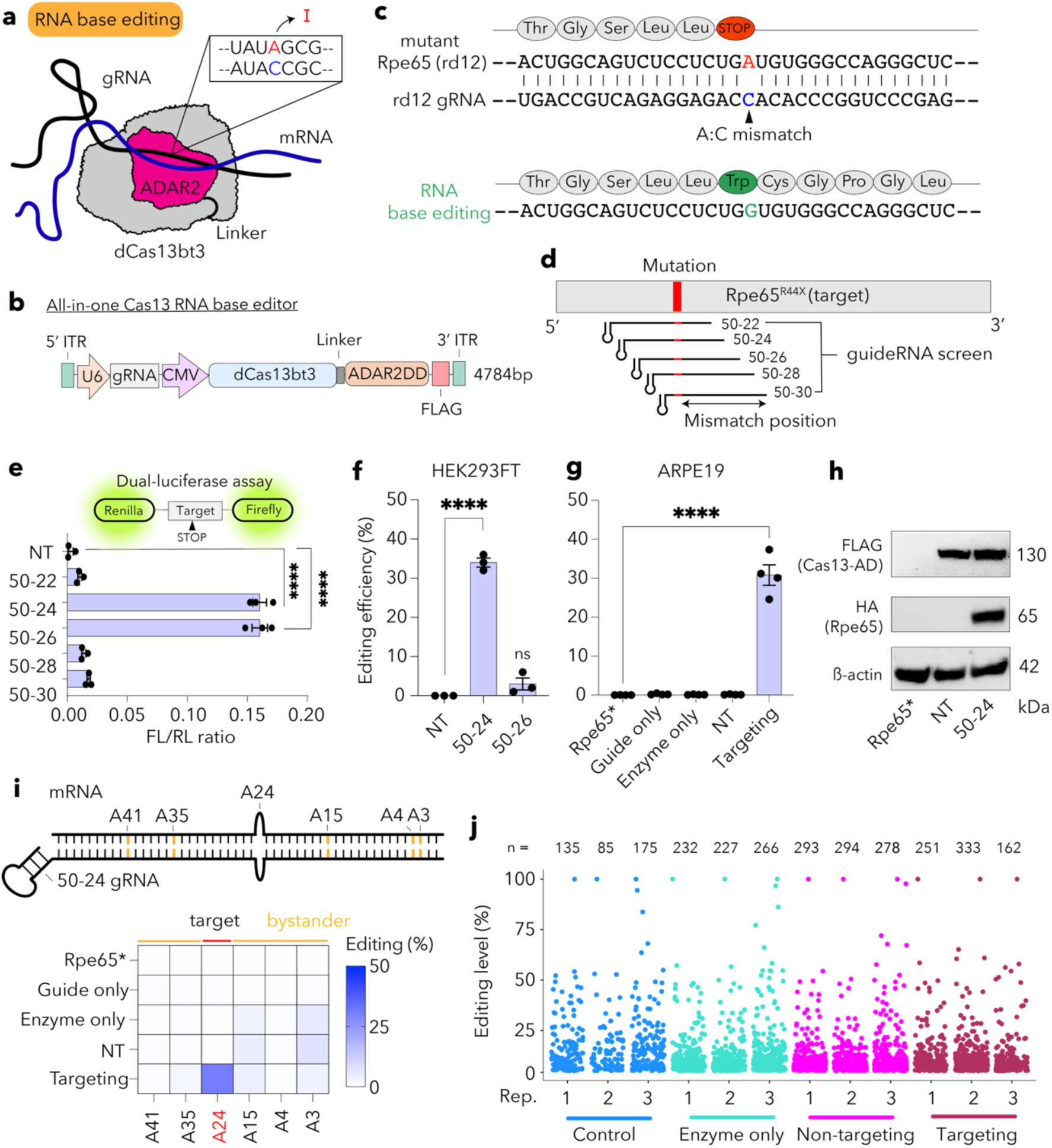
*In vitro* validation of Cas13bt3-ADAR2DD base editor in HEK293FT and ARPE19 cells. **(a)** Mechanism of action for Cas13-mediated RNA base editing. **(b)** Domain structure of all-in-one AAV plasmid carrying CMV-driven dCas13bt3-ADAR2DD and U6-driven gRNA. **(c)** Sequence of *Rpe65*^R44X^ (rd12) nonsense mutation and targeting gRNA with A:C mismatch at target base. RNA base editing achieves an A→I (G) base change, allowing readthrough of the transcript. **(d)** Schematic of gRNA tiling on the target gene for screening of different A:C mismatch positions. **(e)** Dual-luciferase reporter assay for gRNA screening. FL: firefly luciferase, RL: renilla luciferase. **(f)** Editing efficiency by deep sequencing of screened gRNAs in HEK293FT cells against full-length *Rpe65*. **(g)** Editing efficiency by deep sequencing of 50-24 gRNA against full-length Rpe65 in ARPE19 cells compared to gRNA only, dCas13bt3-ADAR2DD (enzyme) only and non-targeting (NT) controls. **(h)** Western blot showing recovery of Rpe65 protein in ARPE19 cells. **(i)** Bystander editing observed in ARPE19 cells with 50-24 gRNA. **(j)** Number of A-to-I events across each treatment group in ARPE19 cells. Data presented as mean ± SEM. ****p < 0.0001; One-Way ANOVA with Tukey’s multiple comparisons test (e, f, and g).

We then targeted a clinically relevant nonsense mutation found in the mouse *Rpe65* gene (c.130C>T, p.R44X), that causes rapid retinal degeneration in mice (known as rd12)^18^. Specific correction of the adenosine in the UG<u>A</u> stop codon will lead to production of tryptophan chain from the UG<u>G</u> codon and restore protein expression (**Figure 1c)**. A panel of gRNAs tiling the target mutation were designed with the A:C mismatch placed at different positions; an A:C mismatch at the 24^th^ base from the 5’ end of the guide is denoted 50-24 gRNA (**Figure 1d and S3a**). A segment of the target gene (198bp) containing the mutation was flanked on both ends with luciferase reporters, allowing the quick identification of optimal gRNAs through a dual-luciferase assay (**Figure S3b**). Here, the 50-24 and 50-26 gRNAs in the dCas13bt3-ADAR2DD system resulted in significant recovery of the 3’ luciferase reporter (Firefly), indicating efficient correction of the point mutation (**Figure 1e**). With the CIRTS system, only the 50-26 sgRNA showed notable luciferase signals (**Figure S3c**). We then delivered these systems into HEK293FT cells along with the full *Rpe65* gene to determine accurate editing efficiencies that account for the native RNA structure of *Rpe65* (**Figure S3d**). Sanger sequencing revealed that the 50-24 gRNA in the dCas13bt3-ADAR2DD led to significant editing of the target mutation, while the 50-26 gRNA only showed less than 10% editing efficiency (**Figure 1f**). With CIRTS, no editing was observed with the 50-26 gRNA (**Figure S3e**). To determine editing rates in a relevant cell line, we then proceeded with evaluation of the dCas13bt3-ADAR2DD and 50-24 gRNA in human retinal pigment epithelium (ARPE-19) cells. To ensure editing was achieved, the presence of both the 50-24 gRNA and dCas13bt3-ADAR2DD, guide-only (50-24 gRNA only) and enzyme-only (dCas13bt3-ADAR2DD only) controls were compared in ARPE19 cells. Here, amplicon sequencing showed ∼30% editing only when dCas13bt3-ADAR2DD carried a 50-24 gRNA, whereas guide-only and enzyme-only controls showed no observable editing (**Figure 1g**). This corresponded to the recovery of the Rpe65 protein, where only the targeting 50-24 gRNA showed recovery of Rpe65 protein, while the mutant *Rpe65* only and non-targeting (NT) conditions showed no protein recovery (**Figure 1h**). This indicated that ∼30% on-target editing was sufficient for effective protein recovery in retinal cells.

We then examined bystander editing of adjacent adenosines in the gRNA binding region, where all conditions with an overexpressed ADAR2DD showed bystander editing at certain sites. For instance, editing rates of ∼3-7% were observed at sites A3 and A15 in enzyme-only, NT, and targeting conditions, whereas guide-only and mutant *Rpe65*-only conditions showed less than 1% editing at these sites (**Figure 1i**). This indicates that the overexpression of ADAR2DD elevates deamination at nearby sites. To further identify the impact of bystander editing and determine specificity of the dCas13bt3-ADAR2DD system, we studied ‘precisely’ edited reads (where only the target base is modified among all edited reads) and found that 79% of reads were precisely edited, indicating that the RNA base editor was highly specific when editing the *Rpe65*^R44X^ mutation (**Figure S4a**). Further investigation into the specific bystander edits also revealed that 90.6% of reads resembled the wild-type (WT) transcript, considering the bystander edits that were synonymous mutations (**Figure S4b**).

To determine deamination rates across the transcriptome, we performed RNA sequencing and identified transcriptome-wide A-to-I editing events. Here, we utilised the recently described DEMINING platform^19^, which leverages the context of variants to identify high-confidence RNA edits. Compared to guide-only control, only the NT and 50-24 gRNA led to a significant increase in total A-to-I editing events, although this effect was more consistently observed with the NT gRNA (**Figure 1j and S4c**). All groups that overexpressed ADAR2DD showed an increased number of events, indicating that exogenous expression of ADAR elevated transcriptome-wide events, as has been shown in several other studies^12, 15, 20^. A-to-I editing events were also marked at chromosomes 1, 7, 17 and 19 (**Figure S4d**), indicating a hotspot for RNA editing in the region. This corresponds to previous studies showing that chromosome 19 possesses the highest editing frequency, accounting for chromosome size^21^. The effect observed with the NT gRNA was notable and prompted us to evaluate if NT gRNAs will especially increase transcriptome-wide events due to unspecific binding during the target search process. We obtained another non-targeting gRNA (NT2), confirmed to have no binding targets in the human transcriptome, and assessed their transcriptome-wide editing profile in a similar manner. Here, we observed that while none of the groups had significantly increased A-to-I editing events, the NT2 gRNA had the highest number of events (**Figure S5a-b**). While inconclusive, the effect of NT gRNAs on transcriptome-wide editing warrants further study to inform on the importance of effective targeting gRNAs. Our results demonstrate that targeted RNA base editing can be achieved in a relevant retinal cell line with high specificity, showing minimal bystander or transcriptome-wide off-target effects.

### Different pathogenic RPE65 nonsense mutations can be corrected by RNA base editing

We then sought to demonstrate RNA base editing efficiency against clinically reported IRD mutations to evaluate its clinical potential. First, all pathogenic G>A nonsense mutations in the human *RPE65* gene were retrieved from the Leiden Open Variation Database (LOVD). These mutations (amounting to a total of 6) were namely c.992G>A, c.993G>A, c.1205G>A, c.1374G>A, c.1379G>A, c.1380G>A (**Figure 2a**) and occurred across the codon preferences for ADAR2 enzymes (**Figure 2b**). In addition to determining the clinical utility of the dCas13bt3-ADAR2DD base editor, targeting these mutations would enable us to identify whether the context of the target base (5’ and 3’ flanking bases) influences editing efficiencies.

**Figure 2.**
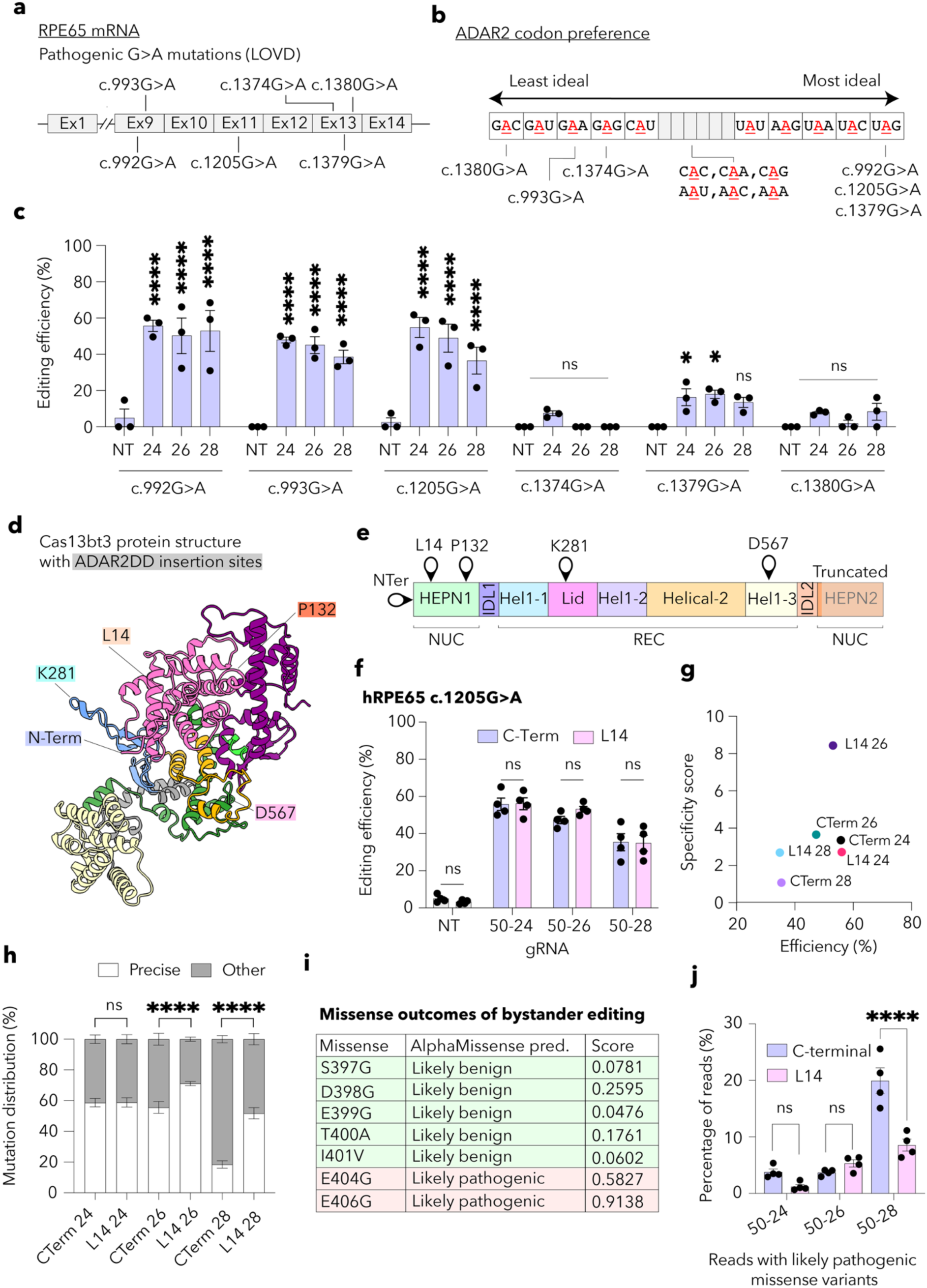
Correction of clinically reported *RPE65* pathogenic G>A nonsense mutations. **(a)** Schematic of *RPE65* mRNA with clinically reported pathogenic G>A mutations in the Leiden open Variation Database. **(b)** ADAR2 codon preference table showing where codon context of each *RPE65* mutation. **(c)** Editing efficiency observed from targeting *RPE65* mutations with non-targeting (NT), 50-24, 50-26 and 50-28 gRNAs, determined from amplicon sequencing. **(d-e)** dCas13bt3 protein and domain structure with potential ADAR2DD insertion sites. NUC: nuclease lobe, REC: recognition lobe. **(f)** Editing efficiency observed with C-terminal (C-Term) and L14 base editors targeting c.1205G>A mutation. **(g)** Specificity scores of the C-terminal and L14 base editor with each gRNA against their on-target editing efficiencies. **(h)** Percentage of precisely edited transcripts with C-terminal and L14 base editing with each gRNA. **(i)** Table of potential missense variants in the binding region of gRNAs with their AlphaMissense pathogenicity predictions and score. **(j)** Percentage of reads with likely pathogenic variants as predicted by AlphaMissense for the C-terminal and L14 base editors with each gRNA. Data presented as mean ± SEM. ∗p < 0.05, ****p < 0.0001, ns: not significant; One-Way ANOVA with Tukey’s multiple comparisons test (c, f, h and j).

Centrally placed A:C mismatches in gRNAs have been found to be efficient by us and others^15^. We therefore designed a panel of three gRNAs (50-24, 50-26 and 50-28) against each of the six mutations to identify editing efficiencies at these sites. Here, we observed high editing rates with the c.992G>A, c.993G>A and c.1205G>A mutations ranging 40% - 60%, while little or no editing was observed with the c.1374G>A, c.1379G>A, c.1380G>A, ranging 0% - 20% (**Figure 2c and S6a**). Firstly, we noted that the preferred U<u>A</u>G codon^22^ occurring in c.1379G>A was not efficiently edited while the less favourable G<u>A</u>A codon occurring in c.993G>A was efficiently edited, indicating factors other than ADAR2 preference alone play a role in RNA editing. Still, the U<u>A</u>G in the c.1379G>A mutation still showed to be more permissive to editing compared to the G<u>A</u>G containing c.1374G>A or G<u>A</u>C containing c.1380G>A mutations. The proximity of the c.1374G>A, c.1379G>A and c.1380G>A mutations indicated that RNA structure at this region might inhibit Cas13-gRNA-mRNA complex formation for efficient editing. Using RNAFold, we predicted the RNA secondary structure of full-length *RPE65* and identified the regions containing the mutations, but no notable differences were found. As RNA structure at the target sequences did not explain the difference in editing rates, we then sought to investigate if there were factors that hindered binding to the RNA targets and thereby led to reduced editing efficacies. Specifically, several RNA-binding proteins (RBPs) exist within cells that may interact with mutation carrying regions with high affinity^23^, making them inaccessible to the RNA base editor. We identified RBP motifs in our target region using the gRNA binding region as the input using RBPMap to determine the number of motifs present in each target region across the six mutations (**Figure S6b**). Interestingly, we plotted the on-target editing efficiency of each gRNA against the number of RBP motifs present in the target region, and an inverse relationship was observed (**Figure S6c**). Target regions of the c.1374G>A, c.1379G>A and c.1380G>A mutations had ∼20-30 RBP motifs present, and all presented with less than 20% on-target efficiency while target regions of c.992G>A and c.993G>A mutations had no RBP motifs with above 40% on-target efficiency. These results corresponded to findings in a recent study^24^ and indicated that the presence of RBP motifs in the target region can significantly reduce the efficacy of RNA base editing and must be considered when developing RNA base editing strategies against other targets.

As we observed a notable decrease in on-target efficiency across the panel of gRNAs tested against the c.1205G>A mutation, with the 50-24 gRNA being the most efficient and the 50-28 gRNA being least efficient, we then sought to understand if gRNA structure played a role in reducing on-target efficiency (**Figure 2c**). Secondary RNA structure prediction of the three gRNA revealed an additional stem loop present in the 3’ end of the 50-28 gRNA (**Figure S7a and b**). To evaluate the change in RNA stability provided by the additional RNA stem loop, we compared the minimum free energy of each gRNA structure and found the 50-28 gRNA to possess the highest RNA stability (MFE = -10.2) while the 50-24 gRNA was lowest (MFE = -9.9) (**Figure S7c**). As Cas13 searches the transcriptome for a complementary binding region, the presence of structured RNA in the target mRNA or gRNA increases the time required for optimal binding (known as dwell time^25^), which could result in Cas13 dissociating and abandoning the target site altogether. Lower or no editing will be achieved in these cases, as evidenced by our findings, suggesting that gRNA secondary structure must also be considered when designing RNA base editors.

Next, we evaluated bystander editing at each of the mutations, where a clear relationship between on-target efficiency and bystander editing was observed. The c.992G>A, c.993G>A and c.1205G>A mutations all had significant bystander editing across the gRNA binding region owing to better accessibility and binding at these regions, while bystander editing was sparse with the c.1374G>A, c.1379G>A and c.1380G>A mutations (**Figure S6a**). Given the relationship between on-target and bystander editing, we determined specificity scores (where the on-target editing rates are divided by the highest bystander editing rates) to identify gRNAs and mutation sites that offer a good trade-off between on-target and bystander editing. Here, we found that targeting the c.1205G>A mutation with a 50-24 or 50-26 gRNA was especially efficient with minimal bystander editing (**Figure S6d**). While c.992G>A and c.993G>A mutations had high editing efficiencies, specificity was low due to adjacent adenosine sites being highly edited. While multiple bystander editing sites were observed, we sought to study whether incorporating an A:G mismatch at the most highly edited bystander sites in these mutations could achieve higher specificity. However, incorporating these A:G mutations only led to loss of on-target activity, suggesting a significant loss of binding from the RNA base editor with the additional A:G mismatch (**Figure S8**). Therefore, alternative methods to reduce bystander editing should be considered.

### Domain-inlaid RNA base editors for improving RNA base editing efficacy and specificity

As A:G mismatches were unable to maintain on-target efficacy and bystander editing occurred at multiple sites, we hypothesised that modifying ADAR binding to the target region can modulate bystander editing levels and provide a more specific base editing system. Domain-inlaid base editors have been previously described as achieving more specific and improved editing outcome^26, 27^, although the exact mechanism through which this is achieved is not yet known. Bystander editing with domain-inlaid RNA base editors has also not been extensively investigated.

To develop domain-inlaid RNA base editors, we first identified unstructured regions in the protein structure of dCas13bt3. We hypothesised that the insertion of ADAR2DD into unstructured regions would not hinder the folding of the fusion protein and thereby produce functional domain-inlaid RNA base editors. We identified four such sites in dCas13bt3 (2 in the nuclease and 2 in the recognition domains), namely L14, P132, K281 and D567, indicating the amino acid positions of the start of these regions (**Figure 2d-e**). We inserted the ADAR2DD sequence at these regions flanked on both ends by short glycine-serine (GS) linkers to produce four domain-inlaid RNA base editors with Cas13bt3. In addition, we also obtained N-terminal fused ADAR2DD for comparison with the original C-terminal and domain-inlaid designs. The newly designed RNA base editors were evaluated for their functionality and editing efficiencies using the mutant mCherry reporter. Here, high fluorescence recovery was observed with the N-terminal and L14 base editors, comparable to the original C-terminal base editor (**Figure S9a**). The other domain-inlaid RNA base editors (P132, K281, D567) showed limited recovery similar to the scrambled gRNA, indicating these were not functional. Amplicon sequencing further revealed that the L14 base editor had comparable on-target efficiency to the C-terminal base editor, while the N-terminal base editor achieved modestly lower on-target editing (**Figure S9b**). Evaluation of bystander editing showed that the L14 editing also had higher specificity scores due to reduced editing at several bystander sites, indicating the L14 insertion of ADAR2DD may provide a safer alternative base editor (**Figure S9c-d**).

As the L14 editor could improve specificity while maintaining comparable on-target efficacy, we next evaluated the L14 base editors against the c.1205G>A mutation and compared their overall efficacy and specificity to those of the C-terminal base editor. First, all three gRNAs tested (50-24, 50-26 and 50-28) resulted in comparable on-target efficiencies between the L14 and C-terminal base editors (**Figure 2f**); efficacy was moderately lower when using the 50-28 gRNA with both designs. A modified bystander editing profile was also observed between the L14 and C-terminal base editors, indicating that ADAR binding to the target region was altered. For instance, A33 was only highly edited with the C-terminal editor using the 50-24 gRNA (**Figure S10a**). There were also bystander sites that had reduced editing with the L14 base editor, for instance A30 with the 50-26 gRNA and A32 with the 50-28 gRNA. To understand the overall specificity between the L14 and C-terminal base editors across the gRNAs tested, we similarly obtained specificity scores and found that the L14 base editor with the 50-26 gRNA led to the highest on-target editing, along with the highest specificity score (**Figure 2g**). As specificity scores only account for the high bystander editing rate at the most susceptible bystander site, we further evaluated the edited amplicons to determine precisely edited transcripts. By identifying “precise” transcripts (where only the target base is changed on the edited transcripts), a more comprehensive profile of bystander editing can be obtained, and sites that are more likely to be simultaneously edited can also be found. Here, we found the L14 base editor to significantly increase the percentage of precisely edited transcripts with the 50-26 gRNA compared to the C-terminal base editor. Interestingly, while the 50-28 gRNA showed comparable on-target editing between the C-terminal and L14 base editors, albeit generally lower than the other gRNAs tested, the percentage of precise transcripts increased two-fold with the L14 base editor, indicating higher specificity (**Figure 2h**). To determine whether this increase in specificity has clinical value, we examined the effect of each bystander A>I change in the bystander region and identified changes that result in non-synonymous codon modifications. The pathogenicity of these codon changes was therefore determined using AlphaMissense^28^, a deep learning tool that predicts pathogenicity of missense variants across the proteome. Here, we found that 2 out of the possible 7 missense variants from bystander editing at this region were pathogenic (**Figure 2i**). Having identified bystander edits that are likely pathogenic, we then determined the percentage of reads post the base editing event that had pathogenic bystander editing. Here, we found that with the 50-28 gRNA, compared to the C-terminal base editor, the L14 base editor significantly reduced the percentage of reads with a pathogenic bystander edit (**Figure 2j and S10b**), suggesting that the domain-inlaid RNA base editor could limit the editing window and allow safer base editing where pathogenic missense variants could occur.

To further explore the utility of the L14 base editor, we similarly targeted the c.992G>A mutation with the L14 base editor and compared the outcomes to those of the C-terminal editor. Here, comparable on-target editing rates were observed between the L14 and C-terminal base editors, again with the L14 base editor performing moderately better with the 50-24 and 50-26 gRNAs (**Figure S11a**). The bystander adenosine profile of c.992G>A mutation was notable for the presence of an adenosine trinucleotide (AAA) immediately adjacent to the target codon. Evidently, adenosines in this codon were hyperedited though in a specific pattern where the second and third adenosines were prone to be hyperedited while the first adenosine showed lower editing (**Figure S11b**), indicating the position of adenosine in the bystander region plays a role in their tendency to be edited. This pattern was however altered when the 50-26 gRNA was used with the L14 base editor, where the first adenosine was edited at a much higher rate compared to the C-terminal base editor (28.95% vs 7.58%, respectively). This suggested that the editing window had shifted with the L14 base editor in a manner allowing editing of the first adenosine. Other notable altered editing sites were also seen, for instance, A12 with the 50-24 gRNA was reduced to ∼6.72% from 17.80% with the L14 base editor and A7 with the 50-28 gRNA was reduced to 4.14% from 15.79% with the L14 base editor. To gain a better understanding in change of specificity between the C-terminal and L14 base editors, we again determined percentage of precise reads between the editors across the panel of gRNAs. Here, only the 50-24 gRNA was found to increase overall precise reads (**Figure S11c**), indicating the L14 base editor with the 50-24 gRNA could yield safer editing outcomes. We similarly identified missense variants and their pathogenicity using AlphaMissense, finding that 4 out of the 7 possible missense variants were pathogenic (**Figure S11d**). With the L14 base editor and the 50-24 gRNA, reads with pathogenic missense reads were also found to be 2-fold less than with the C-terminal base editor (**Figure S11e**), presenting another case where the domain-inlaid editor could improve overall editing outcomes by reducing bystander editing.

### Targeting the clinically significant and large gene, USH2A, untreatable by current gene therapy methods

As the RNA base editor proved effective at editing multiple mutations across the *RPE65* gene, we sought to demonstrate its use against a large gene that cannot be delivered by AAVs, and is, in principle, untreatable by current gene therapy methods. This would show the utility of the RNA base editor in the clinical setting and provide an alternative strategy for treating currently untreatable genetic diseases. To this end, we identified the largest genes with the most reported incidences for IRDs in the LOVD database to determine clinically significant IRDs. Here, among the largest genes with high incidence numbers globally was *USH2A*, which causes Usher syndrome, a combined vision and hearing loss disease (**Figure 3a**). The most common G>A pathogenic nonsense mutation in *USH2A* was c.11864G>A (W3955X), which is also present in the orthologous mouse *Ush2a* as c.11840G>A (W3947X) (**Figure 3b**). As a mouse model with this variant has recently been reported^29^, we designed a panel of gRNAs (50-22, 50-24, 50-26, 50-28, and 50-30) to target the mouse *Ush2a* (**Figure S12a**). As the full *Ush2a* gene would be challenging to deliver into cells using common transfection methods, we incorporated a short gene segment (198bp) from *Ush2a* carrying the target mutation between luciferase reporters (Renilla and Firefly; **Figure S12a**). This would allow us to identify efficient gRNAs using a quick luciferase assay. Here, we found that the 50-26, 50-28, and 50-30 gRNA led to a significant recovery of Firefly luciferase (**Figure S12b**), indicating efficient base editing rates from these gRNAs. We further evaluated these gRNAs using amplicon sequencing to determine editing rates and found that the 50-26 and 50-28 gRNAs were edited at rates of up to 60%, while the 50-30 gRNA was edited at a moderately lower rate of ∼40% (**Figure 3c**). Specificity scores showed that the 50-26 gRNA yielded the highest on-target editing with minimal bystander editing (**Figure 3d and S12c**). The percentage of precisely edited reads was also the highest with the 50-26 gRNA, making it the ideal candidate for further evaluation. However, the 50-28 gRNA, while achieving comparable on-target editing rates to the 50-26 gRNA, had a significantly edited bystander site at A23, making it less specific. We employed the A:G mismatch at this base to evaluate if the method can be used when minimal bystander sites are present to produce a more specific gRNA (**Figure S12d**). Here, we found that the A:G mismatch at the A23 site specifically reduces bystander editing at the site (33.01% to 5.07%) without affecting on-target activity (**Figure S12e**), suggesting the A:G mismatch is a feasible strategy with the dCas13bt3-ADAR2DD base editor, though its use may be limited depending on sequence context as shown by the loss of on-target with the RPE65 mutations. Interestingly, employing the L14 base editor against the W3947X mutation with the 50-28 gRNA did not yield more specific or efficient editing outcomes (**Figure S12f-g**), indicating multiple strategies may have to be considered when engineering RNA base editors for improved efficiency and specificity. The L14 system also did not reduce transcriptome-wide A>I editing compared to the C-terminal system (**Figure S13a-b**), indicating that overexpression of ADAR2DD is a key factor in global RNA editing.

**Figure 3.**
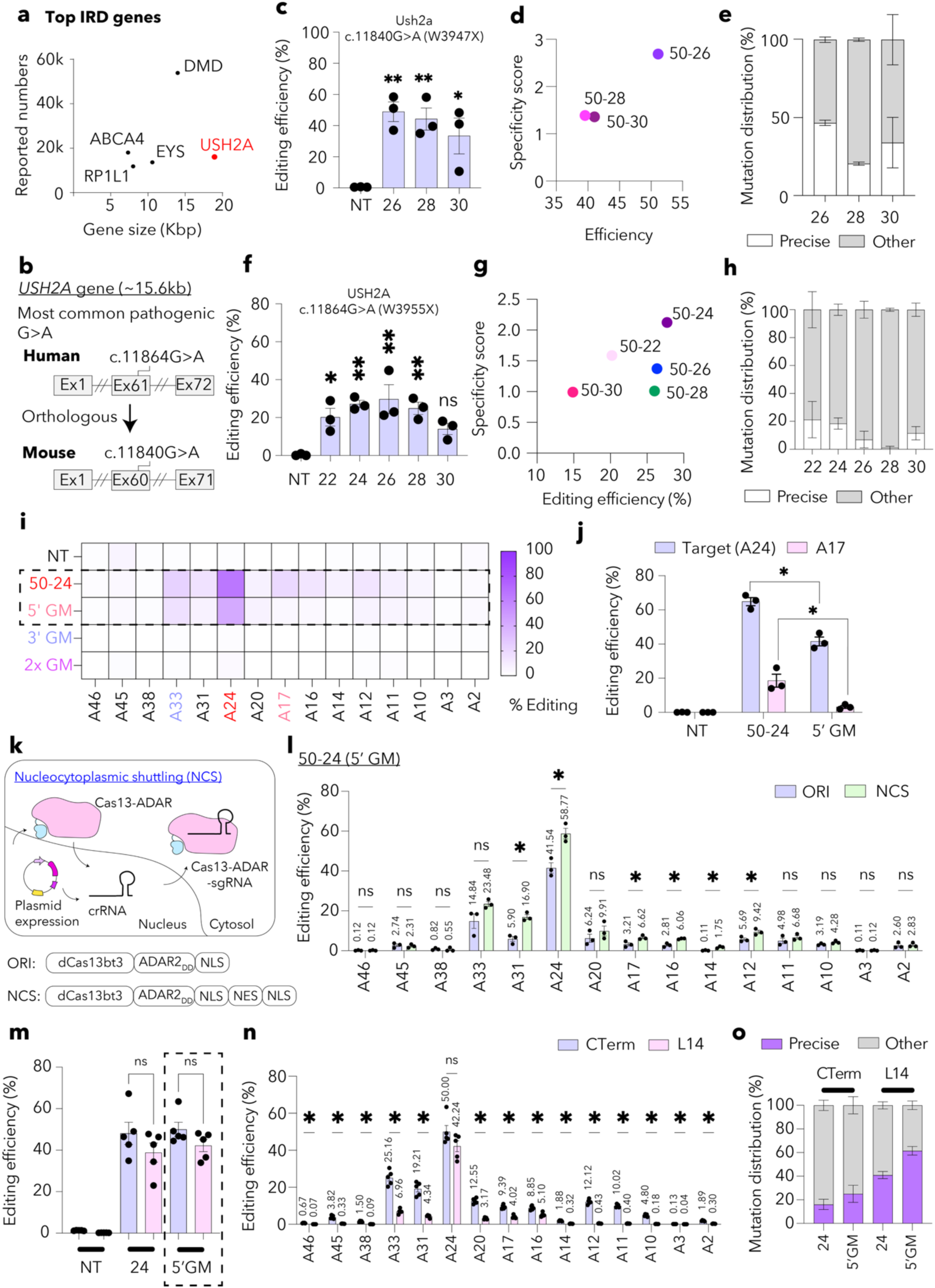
Validation and optimisation of RNA base editing against a large gene, *USH2A* (>15 kb). **(a)** Plot of gene size against reported numbers in the Leiden Open Variation Database (LOVD) showing most prevalent IRD-causing genes that exceed AAV carrying capacity. **(b)** Schematic of locus of most common *USH2A* G>A mutation (c.11864G>A, p.W3955X) and corresponding orthologous mutation in mouse *Ush2a* (c.11840G>A, p.W3947X). **(c)** Editing efficiency determined from amplicon sequencing with a panel of gRNA (non-targeting [NT], 50-26, 50-28 and 50-30) against mouse *Ush2a* c.11840G>A mutation. **(d)** Specificity of the panel of gRNAs against their on-target efficiencies. **(e)** Percentage of precise reads from base editing of *Ush2a* c.11840G>A mutation with the panel of gRNAs. **(f)** Editing efficiency determined from deep sequencing with a panel of gRNAs (NT, 50-22, 50-24, 50-26, 50-28, 50-30) against the *USH2A* c.11864G>A mutation. **(g)** Specificity score of the panel of gRNAs targeting *USH2A* c.11864G>A mutation against their on-target efficiencies. **(h)** Percentage of precise reads from the panel of gRNAs against the *USH2A* c.11864G>A mutation. **(i)** Heatmap showing bystander editing with original 50-24 gRNA, 50-24 gRNA with 5’ G mismatch (GM) or 3’GM or GM at both 5’ and 3’ sites. Bases that were mismatched are indicated in colour, light blue for 5’GM and orange for 3’ GM. The target base is indicated in red. **(j)** A-to-G editing rate at on-target (A24) and bystander site (A17) with 50-24 and 50-24 (5’GM) gRNAs. **(k)** Schematic of nucleocytoplasmic shuttling (NCS) process for dCas13bt3-ADAR2DD base editor (top) and domain structure of original and NCS construct (bottom). **(l)** On-target and bystander editing profile with original and NCS plasmids carrying the 50-24 or 50-24 (5’GM) gRNAs. **(m)** On-target editing efficiency of NCS plasmids with C-terminal or L14 base editors carrying 50-24 or 5’GM gRNAs for targeting *USH2A*^W3955X^. **(n)** Bystander editing profile for C-terminal (C-Term) and L14 base editors with the 5’GM gRNAs. **(o)** Distribution of precise reads after base editing with C-terminal and L14 base editors with 50-24 and 5’GM gRNAs. Data presented as mean ± SEM. ∗p < 0.05, **p < 0.01, ns: not significant; One-Way ANOVA with Tukey’s multiple comparisons test (c, f, j, i, m and n).

Next, we similarly designed a panel of gRNAs for the *USH2A* c.11864G>A mutation to evaluate the dCas13bt3-ADAR2DD base editor for human application (**Figure S14a**). Amplicon sequencing revealed similar editing rates across the panel of gRNAs tested, with only the 50-30 gRNA failing to reach statistically significant editing. The other gRNAs yielded ∼20-40% on-target editing, with the 50-24, 50-26 and 50-28 gRNA achieving the most significant editing rates (**Figure 3f**), with all gRNAs showing bystander editing across the gRNA binding region (**Figure S14b**). Specificity scores did not reveal any gRNA to be significantly more specific than the others, although 50-24 showed the highest score (**Figure 3g**). The percentage of precise reads was notably higher with the 50-22 and 50-24 gRNA compared to the others (**Figure 3h**). We identified missense variants in the gRNA binding region but found none were predicted to be pathogenic (**Figure S14c-d**). As the 50-24 gRNA showed the highest specificity and among the highest on-target activity, we identified significant bystander sites to incorporate A:G mismatches to yield more specific gRNAs. To this end, two sites were chosen (A17 and A33) and gRNA with mismatches in either or both sites were designed to compare editing outcomes with the original 50-24 gRNA (**Figure S14e**). We hypothesised that more gRNA mismatches will ultimately result in a lack of binding of gRNA and target mRNA, and incorporating two gRNA mismatches will allow pushing the A:G mismatch strategy to its limit (**Figure S14f**). We observed that while the 3’GM (A33 mismatched) and 2xGM (both A17 and A33 mismatched) did not yield effective gRNAs, with the 5’GM gRNA (where A17 is mismatched with a G) there was efficient albeit reduced on-target and bystander editing compared to the original 50-24 gRNA (**Figure 3i**), indicating position of the A:G mismatch must be considered employing the A:G mismatch strategy. Amplicon sequencing revealed that bystander editing was almost completely abolished at the A17 site with the 5’GM gRNA, although a significant loss of on-target editing also occurred (**Figure 3j**).

### Nucleocytoplasmic shuttling can improve on-target RNA base editing dCas13bt3-ADAR2DD

As the 5’GM gRNA yielded more specific outcomes compared to the 50-24 gRNA, we postulated that improving on-target editing with the 5’GM gRNA would produce a highly efficient and specific RNA base editing system. As the gRNA is expressed and localised in the nucleus, Cas13 enzymes are often tagged with nuclear localisation signals (NLS) for efficient RNA editing and the lack of a NLS sequence was shown to reduce Cas13 activity^15^. However, protein-coding mRNAs are exported into cytosol where they are more abundant, and therefore initial studies (e.g. REPAIR) had used NES sequences fused to Cas13^16^. Recently, nucleocytoplasmic shuttling was reported as a strategy to improve CasRx knockdown rates by incorporating both NLS and NES sequences in a specific order at the C-terminal end^30^. We sought to investigate nucleocytoplasmic shuttling (NCS) for RNA base editing by incorporating the NCS sequence into the C-terminal end of the dCas13bt3-ADAR2DD fusion protein (**Figure 3k**) and thereby improving on-target editing outcomes with the 5’GM gRNA. First, we observed that incorporating the NCS tail improved on-target editing significantly (41.54% to 58.77%), however this also caused increased bystander editing at most sites (**Figure 3l and S15a**). Significantly increased bystander editing was observed at A31, A17, A16, A14 and A12, emphasizing the tight relationship between on-target and bystander editing (**Figure 3l and S15b**). Despite the increase in on-target editing, the percentage of precisely edited reads remained constant, indicating no improvement in specificity from the NCS sequence (**Figure S15c-d**).

Lastly, to improve final editing outcomes, we incorporated the NCS sequence with the L14 domain-inlaid design and 5’GM gRNA. On-target editing rates were comparable (**Figure 3m**), and surprisingly, bystander editing was significantly reduced at all bystander sites (**Figure 3n**), revealing a highly specific and efficient RNA base editor against the *USH2A* c.11864G>A mutation. We determined the percentage of precisely corrected reads, noting that improvements in precise reads were observed with the 5’GM and L14 designs. Using both strategies together resulted in the highest level of precise reads (**Figure 3o**).

### Demonstrating RNA base editing for recovery of Rpe65 and retinal function in the rd12 mouse

Given the promising editing outcomes and validation of engineering strategies for efficiency and specificity, we sought to demonstrate RNA base editing *in vivo* in the rd12 mouse that presents with early and rapid cone degeneration due to a nonsense mutation (c.130C>T, p.R44*) in the *Rpe65* gene^31^. We packaged the dCas13bt3-ADAR2DD base editor, along with either NT or 50-24 gRNA that we had previously validated *in vitro,* into AAV2/2 vectors (**Figure 4a**). Viral vectors encoding the RNA base editor were co-delivered with AAV2/2-GFP via subretinal injection into the eyes of 4-5-week-old rd12 mice. Retinal pigment epithelial tissues were collected six weeks post-injection for analysis (**Figure 4b**). We first confirmed the distribution of AAV in mouse retinal cells from blue-peak autofluorescence retinal imaging (for GFP expression; **Figure 4c and S16a**) and RT-qPCR amplification of dCas13bt3-ADAR2DD from rd12 RPE cells (**Figures 4d**). Amplicon sequencing from rd12 RPE showed that the 50-24 gRNA could achieve ∼2-4% on-target editing, although this was not consistent across all eyes tested (**Figure 4e**), indicating variable or inefficient virus expression following subretinal injections. To evaluate whether the observed editing led to the preservation of photoreceptor function, we compared cone responses from the rd12 mice with positive editing to those from the NT- and GFP only treated groups. As rod responses may contribute to overall ERG responses, a double-flash ERG was used to isolate cone responses, where the first flash desensitises the rods for a cone-specific response to be recorded from the second flash^32^. Here, we observed increased cone-specific electroretinogram (ERG) responses compared to baseline recordings in the rd12 mice treated with the 50-24 gRNA (**Figure 4g**), and the responses were maintained between baseline and 6 weeks after treatment (**Figure 4f**). This difference was significant compared to both the NT and GFP only groups, indicating that correction of the rd12 nonsense mutation by the RNA base editor led to preservation of cone function (**Figure S17**). No such improvements were observed with the dark-adapted ERG a- or b-wave responses, however, likely owing to the lack of rod photoreceptor responses in rd12 mice. We then performed immunostaining and confirmed that the editing led to recovery of RPE65 protein in the RPEs treated with the 50-24 gRNA only (**Figure 4h and S16b**). To determine if the treatment had any effect on retinal structure, we obtained retinal thickness readings from OCT images and compared them across four regions (2 nasal and 2 temporal) for total retinal and outer retinal thickness (**Figure S18**). Here, while thickness profiles did not differ significantly between the NT and 50-24 gRNA groups for both types of thicknesses, there was a significantly lower outer retinal thickness for both NT and 50-24 groups compared to the GFP group in outer retinal thickness readings (**Figure 4i-j**). As total retinal thicknesses did not significantly vary between the treatment groups, whether this effect is due to toxicity from the dCas13bt3-ADAR2DD delivery or damage from subretinal injections must be further studied. Overall, our data suggest that AAV-mediated subretinal gene delivery of dCas13bt3-ADAR2DD along with 50-24 gRNA enables the correction of the *Rpe65*^rd12^ mutation in mRNA, leading to recovery of Rpe65 protein and partially preserving cone function.

**Figure 4.**
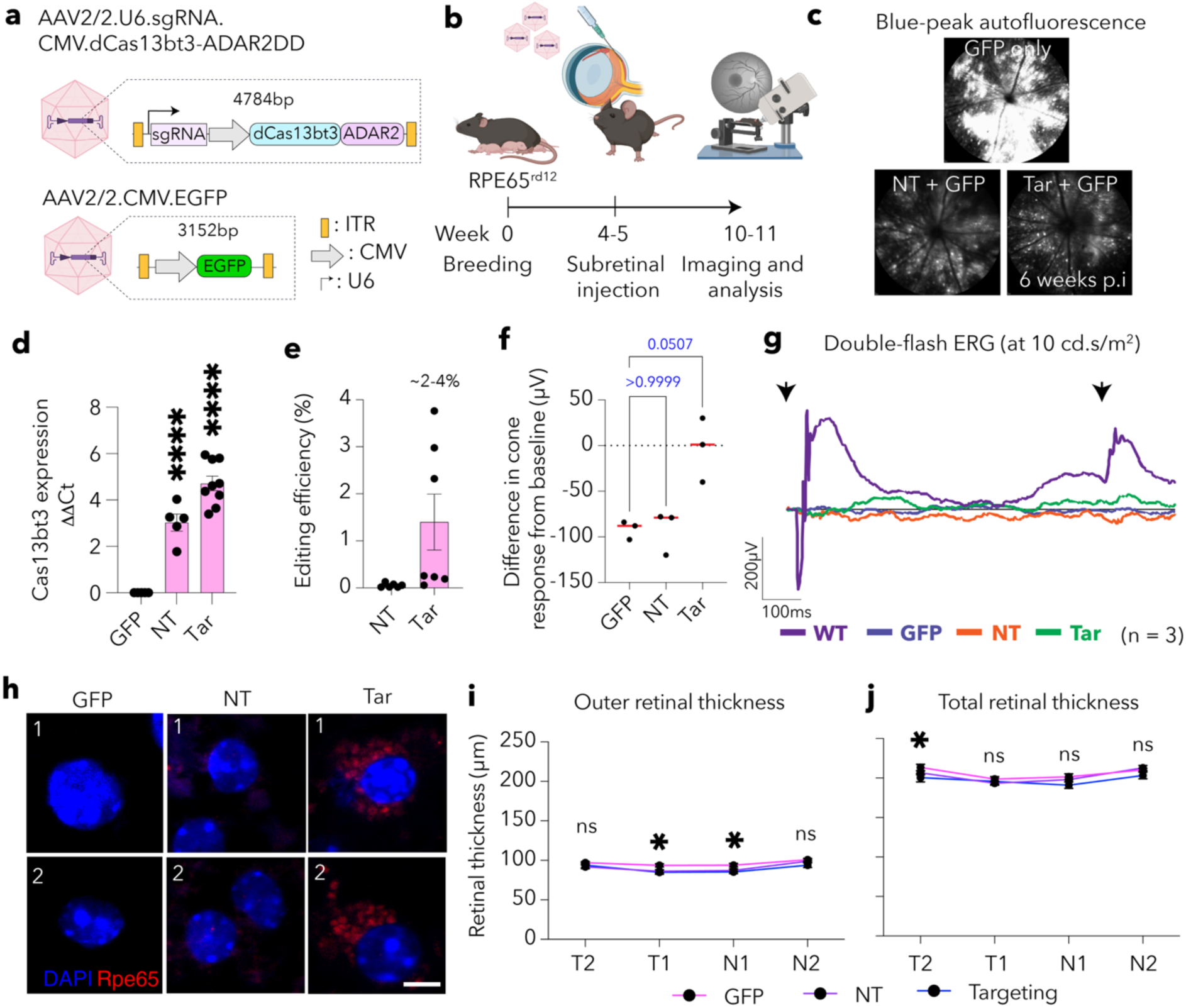
RNA base editing for correction of *Rpe65* gene in *Rpe65*^rd12^ mice. **(a)** Schematic of AAV2/2 virus mix carrying GFP reporter (GFP) and dCas13bt3-ADAR2DD with non-targeting (NT) and targeting (Tar) gRNAs. **(b)** Schematic of experimental procedure for subretinal injection into rd12 mice. **(c)** Blue-peak autofluorescence images taken 6 weeks post-subretinal injection showing expression of GFP reporter. **(d)** RT-qPCR of dCas13bt3-ADAR2DD from the RPEs of rd12 mouse. **(e)** Editing efficiency determined from amplicon sequencing from the rd12 mouse RPEs. **(f)** Difference in cone response from rd12 mice treated with GFP or dCas13bt3-ADAR2DD base editors. **(g)** Representative ERG response from rd12 mice treated with GFP or dCas13bt3-ADAR2DD base editors. Black arrows indicate stimulus **onset**. ERG recordings are averaged from 3 eyes per group. **(h)** Immunofluorescence of rd12 RPE showing expression of Rpe65 protein after treatment with GFP or dCas13bt3-ADAR2DD base editors. Scale bar: 5 µm. **(i-j)** Outer and total retinal thickness of rd12 mice treated with GFP or dCas13bt3-ADAR2DD base editors across two temporal and two nasal regions from the optic nerve centre. Significance values are shown in relation to the GFP treatment group. Data presented as mean ± SEM. ∗p < 0.05, **p < 0.01, ****p < 0.0001, ns: not significant; One-Way ANOVA with Tukey’s multiple comparisons test (d, e, i and j) and Kruskal-Wallis test **(f)**.

### RNA base editing achieves efficient correction and recovery of Ush2a in vivo after intravitreal injection

Next, we obtained the *Ush2a*^W3947X^ mice recently reported for targeting to demonstrate RNA base editing of a large gene *in vivo* for efficacy and safety. As AAV serotypes like AAV2.7m8 have demonstrated expression in photoreceptors from intravitreal delivery, we sought to deliver the base editors similarly to enable a less invasive treatment approach compared to subretinal injections. To this end, we packaged the dCas13bt3-ADAR2DD base editors along with either NT or the 50-26 gRNAs into AAV2.7m8 vectors. As previously done, we also co-delivered AAV2.7m8-GFP reporter to evaluate delivery efficacy (**Figure 5a**). We then delivered the virus mixture via intravitreal injections into 10- to 12-week-old *Ush2a*^W3947X^ mice and assessed editing efficacy and protein recovery using NGS (amplicon sequencing) and immunofluorescence staining 8 weeks post-viral injection (**Figure 5b**). Expression of dCas13bt3-ADAR2DD was first confirmed using RT-qPCR and Western blot. While RNA expression proved to be consistent across the treated eyes (**Figure 5c**), protein expression was either challenging to detect or varied across samples (**Figure S19**). This is likely owing to the differential expression from intravitreal injections. Due to localisation of Ush2a expression in the photoreceptors and overall transduction rate with intravitreal injections, sequencing from the whole retina is likely to underrepresent actual editing rates. To mitigate this effect, we performed dissociation of mice retinae and fluorescence-activated cell sorting with GFP prior to RNA extraction and sequencing (**Figure S20**). Here, we observed 2-20% on-target editing in the eyes treated with the 50-26 gRNA, compared to 0-2% editing in the NT group, indicating that the targeting gRNA could specifically correct the *Ush2a* mutation (**Figure 5d**). To determine if the editing led to recovery of the Ush2a protein, we performed immunostaining for Ush2a in retinal sections and found the retinae treated with the 50-26 gRNA showed expression of Ush2a at the tips of the photoreceptor inner segments unlike retinae treated with the NT gRNA (**Figure 5e**), consistent with localisation of Usherin in mouse photoreceptors to the connecting cilium.

**Figure 5.**
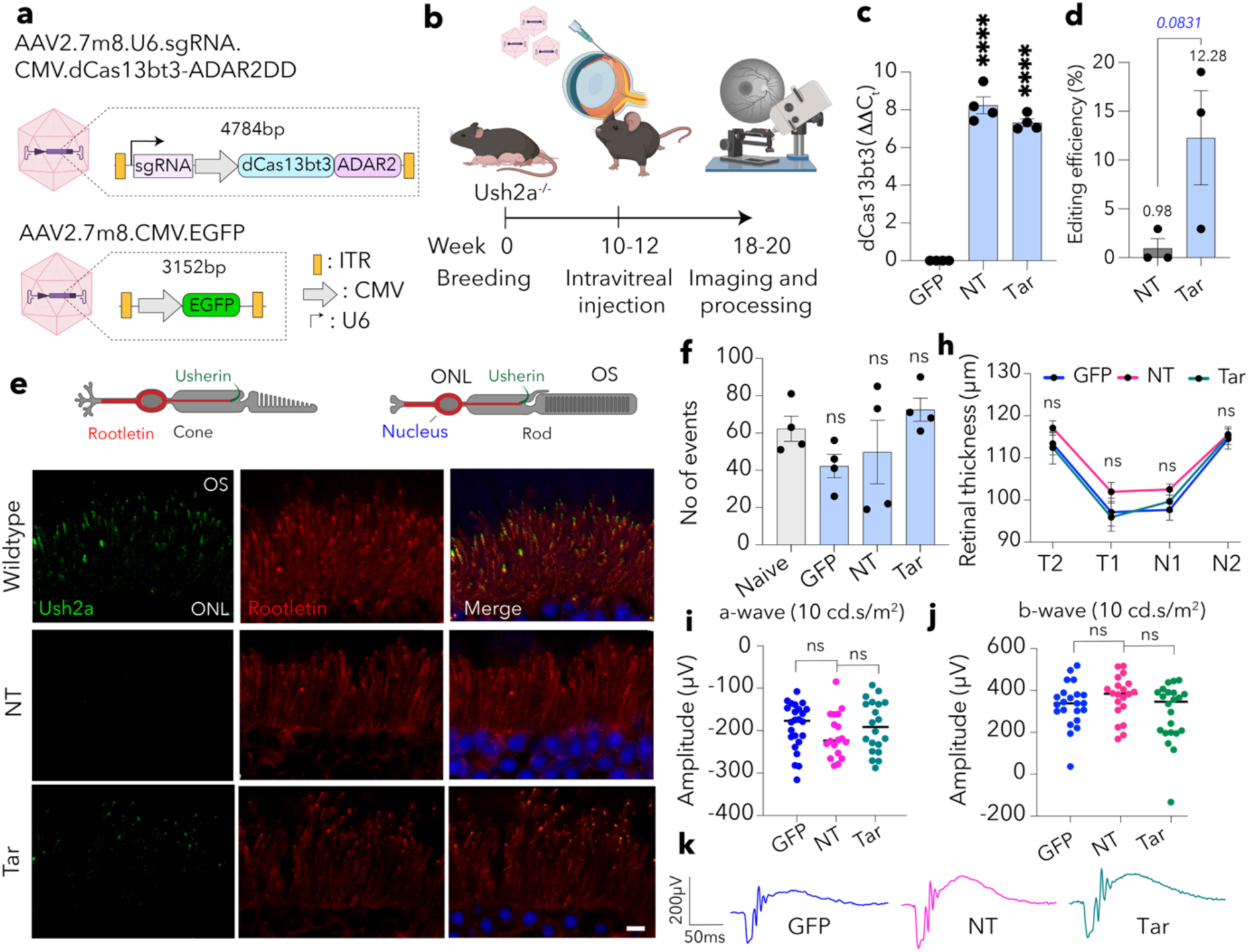
RNA base editing in *Ush2a*^W3947X^ mice. **(a)** Schematic of AAV constructs carrying GFP reporter (GFP) and dCas13bt3-ADAR2DD with non-targeting (NT) and targeting (Tar) gRNAs. **(b)** Experimental procedure for intravitreal delivery into *Ush2a*^W3947X^ mice and endpoint assays. **(c)** RT-qPCR from *Ush2a*^W3947X^ mice retina for expression of dCas13bt3-ADAR2DD. **(d)** Editing efficiency determined from amplicon sequencing from *Ush2a*^W3947X^ mice retina post-dissociation and FACS-sorting. **(e)** Representative immunofluorescence images from *Ush2a*^W3947X^ mice retina showing expression of Usherin protein in the GFP and dCas13bt3-ADAR2DD base editors treatment groups. Schematic of Usherin and rootletin localisation in mice photoreceptors in shown at the top. Scale bar: 5 µm. **(f)** Transcriptome-wide A-to-I events in *Ush2a*^W3947X^ mice retina 8 weeks post AAV injections. **(h)** Retinal thickness determined using OCT images from *Ush2a*^W3947X^ mice. **(i)** A-wave responses and **(j)** b-wave responses (at 10cd-s/ms^2^) from ERG readings 8 weeks after AAV injections. **(k)** Representative ERG traces from GFP and dCas13bt3-ADAR2DD base editors treatment groups showing a-wave responses 8 weeks after AAV injections. Data presented as mean ± SEM. ****p < 0.0001, ns: not significant; One-Way ANOVA with Tukey’s multiple comparisons test (c, d, h, i and j).

As the *Ush2a*^W3947X^ does not present a functional ocular phenotype, we sought to ensure the safety of the virus delivery and RNA base editing therapy with transcriptomic, retinal function and retinal structure analyses. Transcriptome-wide off-target editing rates were determined from RNA sequencing, where no significant differences could be observed between any of the virus-treated groups and the untreated (Naïve) controls (**Figure 5f and S21a-b**). We also identified differentially expressed genes (DEGs) to determine the transcriptomic impact of the virus and the expression of the base editor. First, we did not detect any changes in *Ush2a* transcript counts, indicating the RNA level of *Ush2a* remains constant with the base editing (**Figure S21c**). Differential expression analyses showed that the GFP treatment led to the highest amount of DEGs (6276) compared to the NT and 50-26 gRNA groups (69 and 207, respectively), indicating a strong transcriptomic impact from the reporter virus. This may be due to the stronger expression of this virus in the retina or components of the GFP reporter being genotoxic (**Figure S20d-e**). We then compared outer retinal thickness profiles across the nasal and temporal regions as previously described and found no significant differences between the treatment groups (**Figure 5h**). Dark-adapted visual responses between the treatment groups were also compared from ERG recordings, where again no significant differences were observed in a-wave and b-wave amplitudes, indicating that the virus treatment and base editing therapy did not affect overall retinal integrity/function (**Figure 5i-k**). Together, our data further confirmed that AAV-mediated gene delivery of dCas13bt3-ADAR2DD, along with targeting gRNA, enables the correction of the *Ush2aW*^3947X^ mutation in mRNA, leading to recovery of Usherin protein without causing transcriptome-wide editing or impacting retinal function.

## DISCUSSION

Conventional DNA base editors continue to be hampered by either a lack of specificity, effective delivery or strict PAM requirements. RNA base editing has emerged and shown promise in addressing this drawback, and several approaches using either exogenously or endogenously delivered ADAR enzymes have been described^11, 33, 34, 35, 36^. Here, we adopted the Cas13-fused ADAR2DD and successfully demonstrate methods to optimise gRNA design, engineer a domain-inlaid base editor to improve specificity, and showcase nucleocytoplasmic shuttling constructs to enhance on-target editing. We packaged this in a single AAV vector and demonstrated its potential for therapeutic editing in the retina for inherited retinal disease.

Several studies investigating RNA base editing have shown methods to achieve high on-target efficacy and reduce bystander editing. However, most studies still do not address that clinically occurring mutations do not always exist in a U<u>A</u>G codon context, and mutations that occur in different contexts are likely to be less efficiently edited. Here, we assayed the activity of the dCas13bt3-ADAR2DD base editor across pathogenic G>A nonsense mutations in an entire gene (i.e., *RPE65*) to determine its versatility in addressing clinically relevant mutations. Revealing context preferences with the dCas13bt3-ADAR2DD system enables the tailoring of its use against amenable mutations, while also encouraging the development of alternative methods for editing less amenable mutations. A toolkit of RNA editing methods can thus be identified from which scientists can develop base editing therapies across the mutation landscape.

Despite its advantages, the key concern with RNA editing for gene therapy applications is the potential for undesired editing outcomes on the transcriptome, which we have termed bystander and off-target editing here. Carefully studying these effects will be instrumental in the development of RNA editing gene therapies. In addition to studying transcriptome-wide editing events from RNA sequencing, we have comprehensively studied bystander editing in the gRNA binding region in this work. To this end, we have endeavoured to determine precisely edited reads from on-target editing outcomes, study the impact of each missense variant in the bystander region, determine specificity scores for ranking trade-off between on-target and bystander editing and investigate the incorporation of domain-inlaid editors and A:G mismatches in gRNA based on bystander editing information. These methods not only enable the advancement of base editing therapies in a thoroughly considered manner but also inform future development efforts. Further studies should include detailed analyses of undesired editing outcomes in more relevant models to more accurately determine the translatability of the RNA base editors.

With delivery being a long-standing bottleneck to gene therapy applications, here we have endeavoured to demonstrate the use of RNA base editing with subretinal and intravitreal injections, the two most common methods for retinal delivery. This showcases the ability of the tool to address a wide range of IRD-causing genes that may occur in the inner or outer retina. As DNA editing methods are often preferred for chronic conditions like IRDs, the large size of DNA base editors necessitates the use of alternative delivery vectors such as virus-like particles, lentiviruses, nanoparticles or dual-AAV approaches, which have been described in recent studies^37, 38, 39^. While precise and permanent correction of the mutation in the target cells has been shown, the extent of delivery and level of efficacy remain suboptimal. Our approach promises to overcome the drawback of a lack of delivery and editing efficacy. Delivery with AAVs also indicates potential for lasting therapeutic benefit. As the actual longevity of AAV expression is not fully understood, the possibility of redosing must be considered with an AAV-mediated RNA base editing therapy. To this end, orthogonal AAVs have been proposed and studies with Luxturna have shown potential for redosing AAV therapies^40, 41^. Follow-up studies with currently ongoing AAV gene therapy trials will inform on the durability of therapeutic benefit and potential mitigation strategies.

A major goal of this work was to enable gene therapy for IRDs caused by large genes, where traditional methods like AAV-mediated delivery (e.g., Luxturna) are insufficient. DNA base editors have largely not been able to overcome this challenge due to the large size of SpCas9, PAM requirements and efficacy of alternative Cas9 or Cas12 enzymes. We have demonstrated here that the dCas13bt3-ADAR2DD base editor does not require PAMs, allowing for targeting across the transcriptome, is highly efficient across various contexts, and can address mutations in large genes and prevalent mutations. Mutations in large genes remain largely untreatable and the dCas13bt3-ADAR2DD tool offers a new tool for common diseases with large genes such as Usher syndrome and Stargardt disease. Importantly, the *in vivo* efficacy of ∼12% via intravitreal injection demonstrated in the *Ush2a*^W3947X^ model in this study is notable, and further optimisation to achieve specific expression in photoreceptors has the potential to offer a new gene therapy for Usher syndrome.

Several RNA base editing studies have been recently described for various modifications and improvements to RNA base editing specificity and efficacy^42, 43, 44^. With the base editors having been extensively optimised to achieve high on-target activity with minimal or no bystander editing, a clinical translation pathway must be realised and demonstrated with testing in appropriate human-relevant models for therapeutic application. To this end, we have demonstrated the dCas13bt3-ADAR2DD tool in two different mouse models using two distinct delivery methods; however, further work on human-relevant mutations would strengthen the clinical applicability of these results. For instance, embryonic or iPSC-derived organoid models carrying the human mutations of interest can be targeted with the gRNAs identified from our *in vitro* work to determine efficacy and specificity in a human model. As our *in vitro* work utilised overexpression systems for the target mutation, studies in organoid models would reflect accurate editing and bystander rates. The toxicity of base editors can also be studied comprehensively using these models, employing multi-omic approaches to understand the impact on transcriptome-level perturbations, genotoxicity, and protein expression changes. The impact on the development of retinal structures will also help determine appropriate treatment timepoints for gene therapy. These findings would be broadly beneficial to the retinal gene therapy community working on similar tools for treating retinal disease.

## AUTHOR CONTRIBUTIONS

S.K. and G-S.L. conceived the study. S.K., H-A.C., D.A., A.B., L.H. and C.D.L. performed experiments. S.K. and Y-W.H. performed computational analyses. S.K., C.D.L., A.G-C. and G-S.L. designed experiments and analysed the results. J.Y., A.W.H., F.L., L.E.F., L.S.C., and A.G-C provided critical resources. F.L., L.S.C. and G-S.L. acquired funding. A.G-C. and G-S.L. supervised the research. S.K. wrote the manuscript with input from all authors.

## ACKNOWLEDGMENTS

The authors wish to thank Gavin Knott for assistance with AlphaFold2 prediction in designing domain-inlaid RNA base editors, Paula Fuller-Carter for managing the *Rpe65^Rd12^*mouse, and Anh Vu Truong for helping with NGS sample preparation. The authors also extend their gratitude to the Walter and Eliza Hall Institute of Medical Research for producing the *USH2A*^W3947X^ mouse, VectorBuilder for producing AAV vectors, Azenta Life Sciences, and the Australian Genome Research Facility for conducting RNA sequencing and Sanger Sequencing. Schematics in figures were designed using Biorender.com.

## DATA AND MATERIALS AVAILABILITY

RNA sequencing data generated from this study will be deposited in GEO and SRA. Other data generated during and/or analysed during the current study will be available from the corresponding author on reasonable request. Supporting data and source data can also be found in the supplementary files.

## MATERIALS AND METHODS

| Name | Manufacturer | Catalogue Number/ Version |
| --- | --- | --- |
| ***General lab reagents*** |  |  |
| 2-YT Broth | Thermo Fisher Scientific | 22712020 |
| NEB® Stable Competent E. coli (High Efficiency) | New England Biolabs | C3040I |
| NEB® 10-beta/Stable Outgrowth Medium | New England Biolabs | B9035S |
| Carbenicillin Disodium Salt | Thermo Fisher Scientific | 10177012 |
| GelRed® nucleic acid gel stain | Biotium | 41003 |
| Phosphate buffered saline (PBS) | Sigma-Aldrich | P3813-10PAK |
| Tween-20 | Sigma-Aldrich | P1379-500ML |
| Triton-X-100 | Amresco | 0694-1L |
| RNA Later | Thermo Fisher Scientific | AM7020 |
| Trimethoprim | Sigma-Aldrich | T7883 |
| RIPA Lysis and Extraction Buffer | Thermo Fisher Scientific | 89900 |
| Pierce™ Phosphatase Inhibitor Mini Tablets | Thermo Scientific™ | A32957 |
| NuPAGE™ MES SDS Running Buffer (20X) | Invitrogen™ | NP0002 |
| NuPAGE™ LDS Sample Buffer (4X) | Invitrogen™ | NP0007 |
| NucBlue™ Live ReadyProbes™ Reagent (Hoechst 33342) | Invitrogen™ | R37605 |
| ***Cell culture*** |  |  |
| Dulbecco's Modified Eagle Medium (DMEM), high glucose | Gibco™ | 11965092 |
| DMEM/F-12 | Gibco™ | 11320033 |
| Foetal Bovine Serum (FBS) | Bovogen | SFBS-AU |
| Antibiotic-Antimycotic (100X) | Gibco™ | 15240062 |
| GlutaMAX™ Supplement | Gibco™ | 35050061 |
| TrypLE™ Express Enzyme (1X), phenol red | Gibco™ | 12605028 |
| Dulbecco's Phosphate-Buffered Saline (PBS) | Gibco™ | 14190144 |
| Dimethyl sulfoxide | Sigma-Aldrich | D4540-1L |
| Lipofectamine™ 3000 Transfection Reagent | Invitrogen™ | L3000001 |
| ***Animal procedures*** |  |  |
| 0.5% Proxymethacaine hydrochloride (Alcaine™) | Alcon® | N/A |
| 0.5% Tropicamide (Mydriacyl) | Alcon® | N/A |
| GenTeal™ Gel | Alcon® | N/A |
| Ketamine, 100 mg/mL | Troy Laboratories | Ketamil |
| Xylazine, 100 mg/mL | Troy Laboratories | Ilium Xylazil-100 |
| **Commercial kits** |  |  |
| RNeasy Mini kit | Qiagen | 74106 |
| Monarch® Total RNA Miniprep Kit | New England Biolabs | T2010S |
| High-Capacity cDNA Reverse Transcription Kit | Thermo Fisher Scientific | 4368814 |
| Q5® Hot Start High-Fidelity Master Mix | New England Biolabs | M0494S |
| TaqMan™ Fast Advanced Master Mix | Thermo Fisher Scientific | 4444557 |
| SYBR™ Green Universal Master Mix | Thermo Fisher Scientific | 4309155 |
| Dual-Luciferase® Reporter Assay System | Promega | E1910 |
| **Antibodies** |  |  |
| Anti-rootletin (IF 1:200) | Merck | ABN1686 |
| Anti-Ush2a (IF 1:1000) | A gift from Jun Yang | N/A |
| DYKDDDDK Tag Antibody (IF 1:500, WB 1:1000) | Cell Signalling Technology | #2368 |
| HA-Tag (C29F4) Rabbit mAb (WB 1:1000) | Cell Signalling Technology | #3724 |
| Anti-RPE65 antibody [EPR22579-44] | Abcam | ab231782 |
| Anti-NucBlue™ Live ReadyProbes™ Reagent (Hoechst 33342) | Invitrogen™ | R37605 |
| Donkey anti-Rat IgG (H+L) Highly Cross-Adsorbed Secondary Antibody, Alexa Fluor™ 594 | Invitrogen™ | A-21209 |
| Goat anti-Rabbit IgG (H+L) Cross-Adsorbed Secondary Antibody, Alexa Fluor™ 568 | Invitrogen™ | A-11011 |
| Goat anti-Chicken IgY (H+L) Cross-Adsorbed Secondary Antibody, Alexa Fluor™ Plus 647 | Invitrogen™ | A32933 |
| Goat anti-Chicken IgY (H+L) Secondary Antibody, Alexa Fluor™ 488 | Invitrogen™ | A11039 |
| **Equipment** |  |  |
| HRA + OCT Spectralis | Heidelberg Engineering | N/A |
| Leica Stellaris 5 confocal microscope | Leica Microsystems | N/A |
| ColourDome Stimulator | Diagnosys LLC | N/A |
| **Software** |  |  |
| Prism 10 | GraphPad Software Inc. | 10.1.1 |
| ZEN 2 Pro | ZEISS | 3.6 |
| Illustrator | Adobe Inc. | 28 |
| Photoshop | Adobe Inc. | 25 |
| Image Lab | Bio-Rad | 6.0 |
| BioRender | BioRender | N/A |
| Chimera X | University of California San Diego | 1.3 |
| Snapgene Viewer | Dotmatics | 7.1.0 |
| Benchling | Benchling Inc. | 10 |
| EditR | University of Minnesota | 1.0.10 |
| RStudio | Posit PBC | 2023.09.1 |
| CRISPResso2 | Broad Institute | 2.2.14 |
| RNAFold | University of Vienna | 2.6.3 |

### Primers used in this study

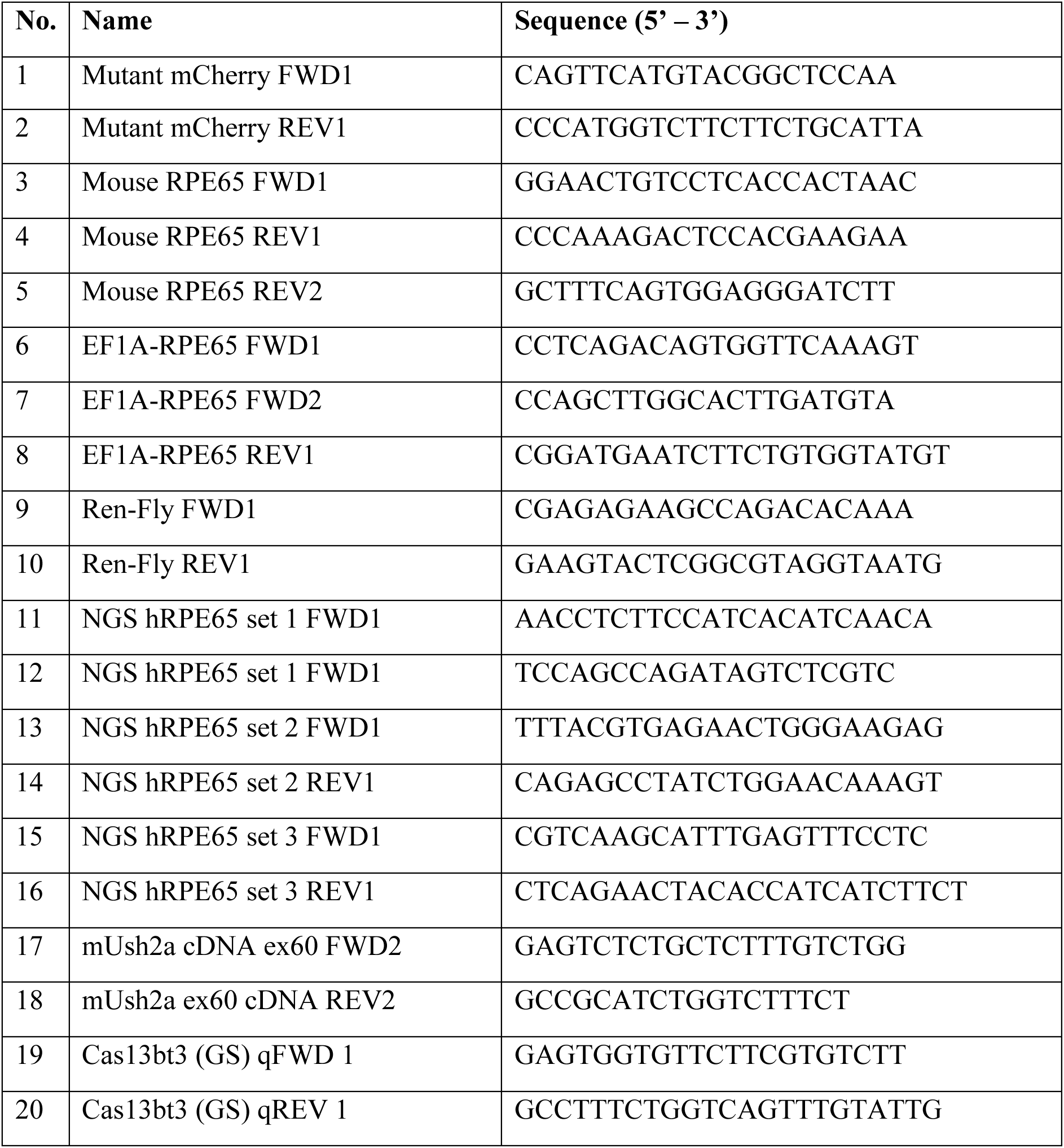

#### Guide RNAs used in this study

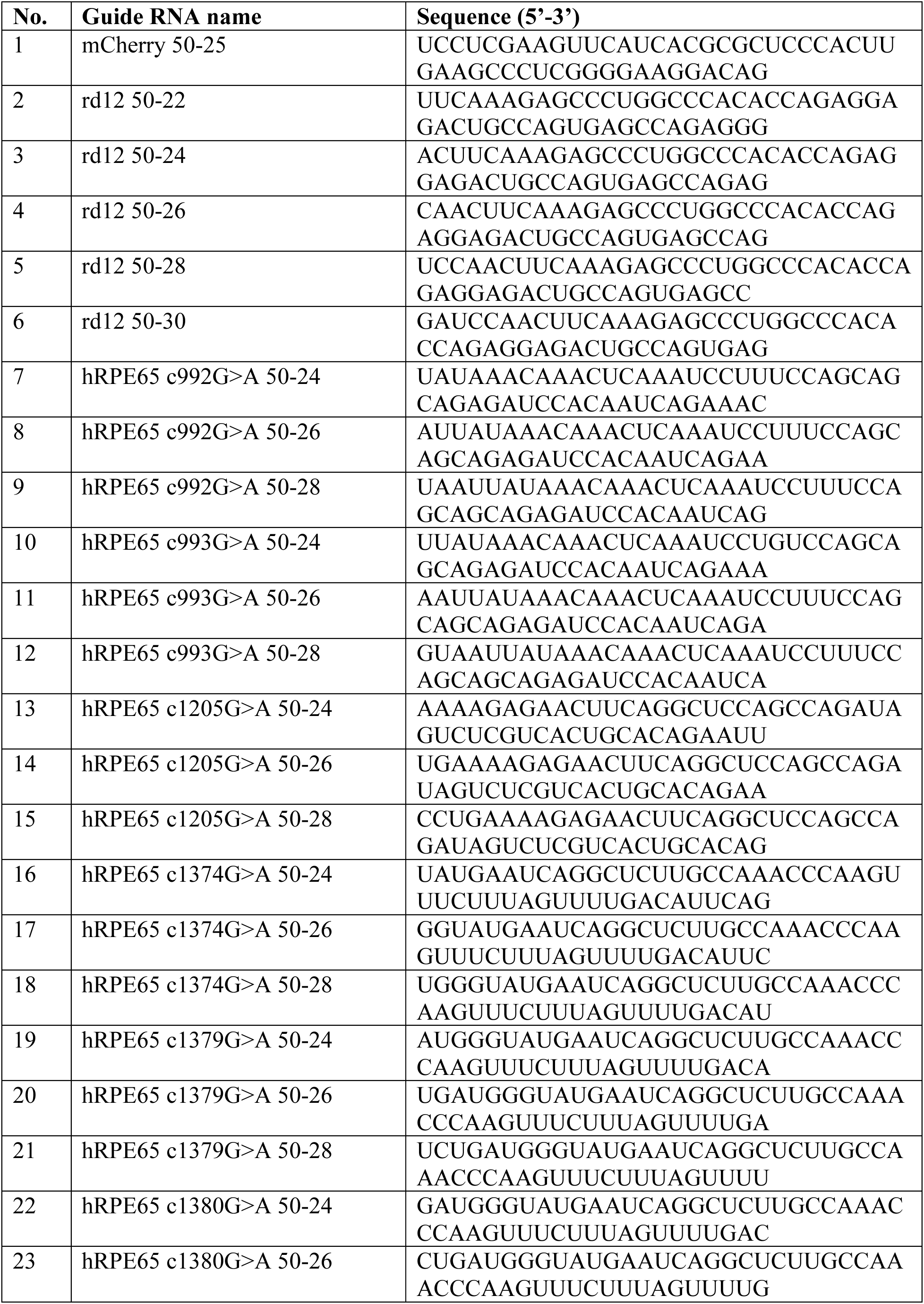
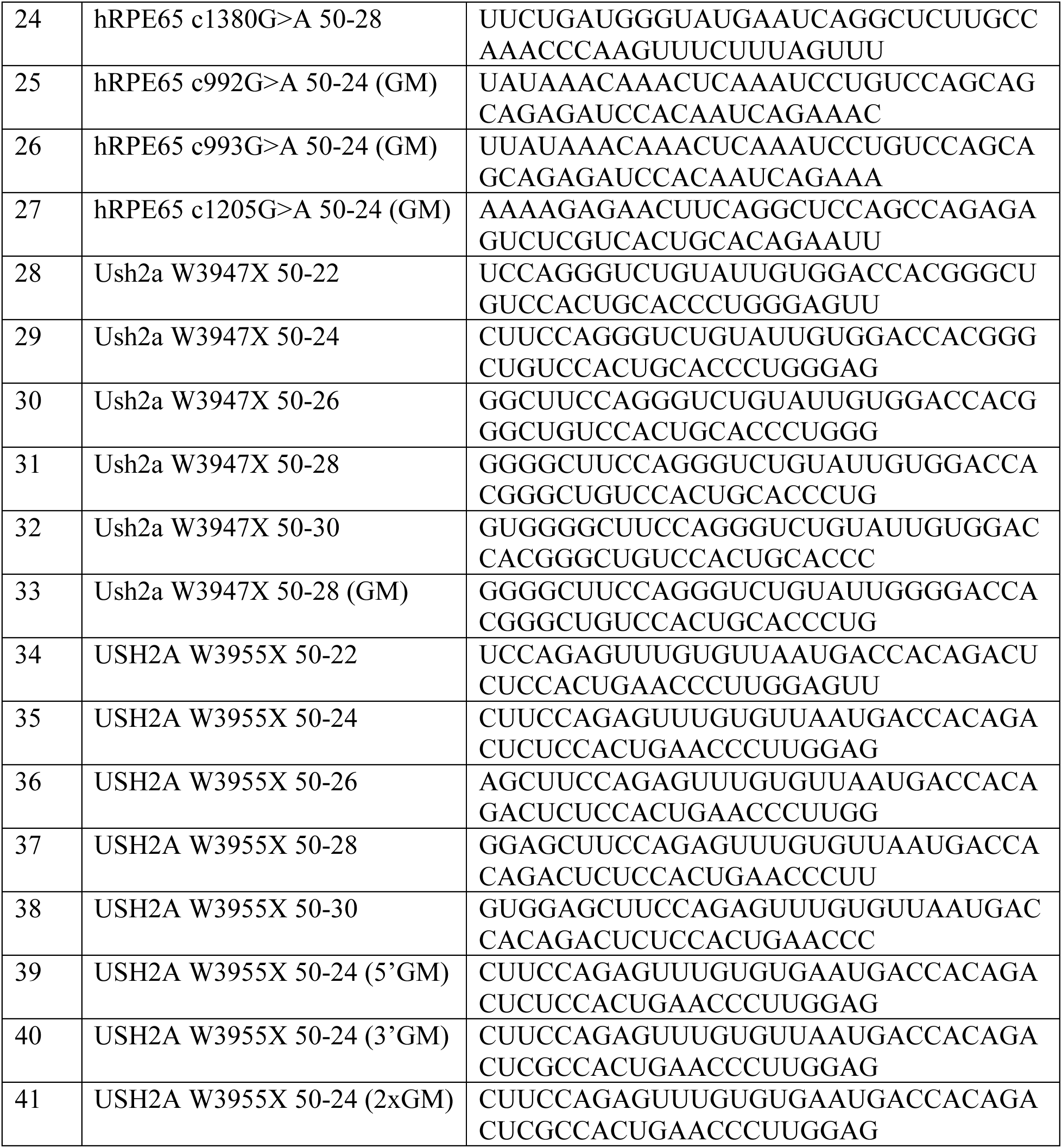

### Cell culture

HEK293FT cells (American Type Culture Collection) were maintained in Dulbecco’s Modified Eagle’s Medium (DMEM; Thermo Fisher Scientific) supplemented with 10% (v/v) fetal bovine serum (FBS, Bovogen Biologicals) and 1% (v/v) antibiotic-antimycotic (Thermo Fisher Scientific) in a humidified 5% CO_2_ incubator at 37°C. ARPE19 (American Type Culture Collection) cells were maintained in DMEM/F-12 supplemented with 10% (v/v) FBS and 1% (v/v) antibiotic-antimycotic.

### Transfection

Transfections were performed using 1:3 Lipofectamine 3000 ratio as per manufacturer’s instructions in 12-well plates. HEK293FT and ARPE19 cells were seeded at 3 × 10^5^ cells per well a day before transfection. Target plasmids (mutant mCherry, mutant human and mouse RPE65 and USH2A, 0.75μg) were co-transfected with base editor constructs (CIRTS or dCas13bt3, 0.75μg). Cells were harvested 48 hours after transfection for luciferase assay or RNA extraction.

### Dual-luciferase assay

Dual Luciferase assay was performed using the Dual-Luciferase^®^ Reporter Assay System (Promega^®^ Corporation) as per the manufacturer’s instructions. Briefly, cells were harvested in 1X DPBS and centrifuged at 16,000g for 2 minutes. The supernatant was discarded, and pellet was resuspended in 250 µL of the provided 1X Passive Lysis Buffer. Cells were then incubated in ice for 10 minutes before centrifugation at 16,000g for 2 minutes. The supernatant was transferred to white opaque 96-well plates for luciferase assay. Two luminescence readings were obtained, one after the addition of LAR II reagent and one after the addition of Stop & Glo substrate.

### RT-PCR

At 48 hours after transfection, RNA was extracted from cells using the Monarch^®^ Total RNA Miniprep Kit and 200ng of RNA was converted to complementary DNA (cDNA) using the High-Capacity cDNA Reverse Transcription Kit (Applied Biosystems™) according to the manufacturers’ instructions. cDNA was diluted with water at a ratio of 1:10. 2μL of cDNA was then used for PCR with respective primers to amplify the target region containing the target mutation. Sanger sequencing of amplicons was performed by the Australian Genome Research Facility (AGRF).

For quantification of dCas13bt3-ADAR2DD RNA from mouse retinae, the retina was dissected from the mouse eyeballs and collected in DNA/RNA Protection reagent (New England Biolabs) and reverse transcribed as previously described. Quantitative PCR was then performed using custom-made primers targeting dCas13bt3 using the SYBR™ Green Universal Master Mix (Thermo Fisher Scientific) using the manufacturers’ instructions.

### Sequencing analyses

Editing efficiencies were determined using Sanger sequencing and amplicon sequencing. PCR was performed with cDNA using Q5 High-Fidelity Polymerase (New England Biolabs) for 30 cycles to obtain 150-350bp amplicons for sequencing. Sanger sequencing was performed by the Australian Genome Research Facility (AGRF), and amplicon sequencing was performed by Azenta Life Sciences. EditR was used to determine editing efficiencies from Sanger sequencing. CRISPResso2 was used to determine on-target and bystander activity from amplicon sequencing. Specificity scores were determined as previously described^24^, using on-target editing over the most highly edited bystander site in the gRNA vinding region. AlphaMissense^28^ predictions for missense variants were obtained from the ProtVar database^45^.

### RNA structure prediction

Secondary structures of RNA sequences were predicted using RNAFold from ViennaRNA package 2.0^46^. As Cas13bt3 processes crRNA to produce a mature crRNA with DR at the 3’ end, the corresponding RNA sequence was input into RNAFold to predict structures for gRNAs. For mutant *RPE65* genes, the entire ORF in the plasmid (start codon to stop codon, containing HA tag) was used as the input for RNAFold.

### A-to-I editing events quantification from RNA sequencing

RNA sequencing (whole transcriptome) was performed by Azenta Life Sciences (150bp paired-end reads). A-to-I editing events were determined using the DEMINING pipeline^19^. Briefly, RNAseq reads were subject to quality control (QC) by FastQC, followed by adaptor trimming by Trimmomatic and alignment to hg38 human genome using HISAT2. Variants were identified using SamTools command mpileup. High-confidence mutations were filtered with the following parameters: base quality ≥ 20, mutation reads ≥ 2, hits per billion mapped bases (HPB) ≥ 3, and mutation frequencies ≥ 0.05. DNA mutations were filtered using co-occurrence mutation context (CMC) prediction from DEMINING output. A-to-G mutations that were classified as RNA mutations were filtered and sites that were supported by at least 100 reads were further filtered to determine total editing events.

### Differential expression analyses

To identify differentially expressed genes, RNA-seq reads were first mapped and quantified using salmon^47^. Differentially expressed genes were then identified using the DESEQ2 package^48^. Genes differentially expressed with >2-fold change and 0.05 p-value were determined to be significant.

### Animals

All animal experiments were approved by the Saint Vincent’s Institute for Medical Research Animal Ethics Committee (AEC) approval number: 019/24, and conducted in accordance with the ARVO Statement for the Use of Animals in Ophthalmic and Vision Research. *Ush2a*^W3947X^ mice were obtained through the introduction of a point mutation of c.11840G>A (W3947X) mutation into the *Ush2a* gene in C57BL/6 by the Melbourne Advanced Genome Editing Centre (MAGEC) at Walter and Eliza Hall Institute of Medical Research (WEHI). The *Rpe65*^rd12^ strain (B6(A)-Rpe65rd12/J) mice were obtained from The Jackson Laboratory.

### Electroretinogram (ERG)

Electroretinogram (ERG) readings were measured immediately prior to subretinal/ intravitreal injections under ketamine-xylazine anaesthesia. Animals were dark-adapted overnight for at least 12 hours before being anesthetized. ERGs were recorded using a gold wire loop on the corneas, with optimal contact between the cornea and the electrode assured using a drop of saline. The reference gold-cup electrode was placed in the mouth, and the ground needle electrode was inserted into the tail. Preparation for ERG recordings was conducted under dim red light. All recordings were performed using an Espion system (Espion e3, Diagnosys LLC; USA). An average of at least three consistent recordings were taken for each stimulus strength. The stimulus strength ranged from -4.0 to 2.0 log cd.s/m^2^. In addition, a double flash recording protocol presented at 10 cd.s/m^2^ with the inter-stimulus interval of 0.8s was performed to obtain an isolated cone response.

### Optical coherence tomography (OCT)

Mice underwent simultaneous spectral domain OCT and confocal scanning laser ophthalmoscopy using the Spectralis HRA+OCT (Heidelberg Engineering, Heidelberg, Germany). A single high-resolution horizontal line scan was obtained after averaging 100 frames using the automatic real-time (ART) mode. A horizontal volume scan (35°x30°) was obtained after averaging 30 frames similarly. Blue-peak autofluorescence was obtained to determine GFP fluorescence.

### Intravitreal injections

At the age of 10-12 weeks, *Ush2a*^W3947X^ mice received intravitreal injections while under ketamine-xylazine anaesthesia, after topical treatment of tropicamide (0.5%) and phenylephrine (2.5%). Intravitreal injections were performed under a surgical microscope using a 33G blunt Hamilton needle mounted on a 10 μL glass syringe (Hamilton Co., Reno, NV) through the pars plana at the temporal side of the eye. The classical transcleral approach with the tip of a needle tangentially inserted through a local sclerotomy was used^49^. Injections were performed slowly and at completion the needle was held in place for at least 10 seconds before removing to minimise the reflux/egression.

### Subretinal injections

At the age of 4-5 weeks, rd12 mice were anesthesised with ketamine-xylazine and both eyes were treated with tropicamide (0.5%) and phenylephrine (2.5%) eye drops. Following ERG and OCT measurements, they received a trans-scleral subretinal injection of ∼0.5 μL (2×10^10^ vg/μL) with a 40G blunt needle after a scleral tunnel was created using a 30G needle. The procedure was visualized with an operating microscope through a dilated pupil. Immediately after the injection, the fundus was examined to check for complications and if present, the animal was removed from the study.

### Immunohistochemistry

For retinal flatmounts, mice eyeballs were enucleated and placed in 4% paraformaldehyde (PFA) for 1 hour before the cornea and lens were removed, 4 radial cuts were made towards the optic head and retinae were stored in PBS for subsequent immunostaining in 4°C. For retinal cryosections, eyeballs were enucleated and fixed in 4% PFA for 2 hours prior to dissection to remove cornea and lens. Eyecups were then subject to sucrose gradient: 10% for 2 hours, 20% for 1 hour and 30% overnight in 4°C. Eyecups were then placed in optimal cutting compound and stored at -80°C until cryosectioning. Sections 12 μm thick were obtained for immunostaining. Sections were washed briefly in 1X PBS before blocking with Dako serum-free protein block (Agilent Dako, X0909) for 1 hour in the dark at room temperature. After washing in 1X PBS 3 times, sections were incubated in primary antibody diluted in Dako antibody diluent (Agilent Dako, S0809).

## SUPPLEMENTARY FIGURES

**Supplementary figure 1.**
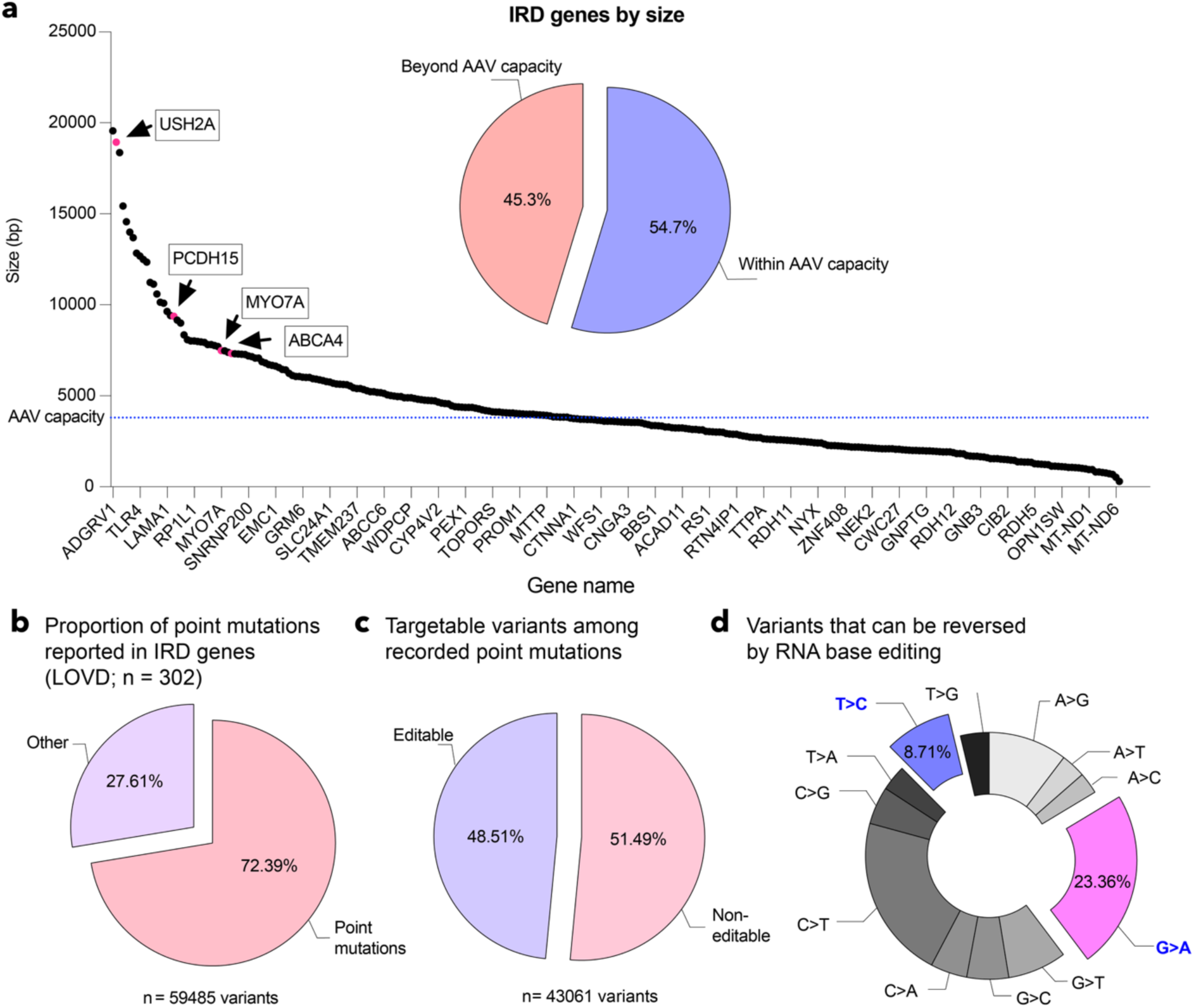
Gene variants causing inherited retinal diseases. **(a)** Plot of IRD-causing genes against their sizes. The blue line on the y-axis indicates practical AAV carrying capacity. The inset pie chart shows the number of genes exceeding the AAV carrying capacity (3.8kb). **(b)** Pie chart showing the number of point mutations among reported variants, **(c)** editable variants among reported point mutations, and **(d)** breakdown of 12 substitution mutations among reported point mutations. Mutations that can be reversed by current RNA base editors, G>A and T>C, are coloured magenta and blue, respectively.

**Supplementary figure 2.**
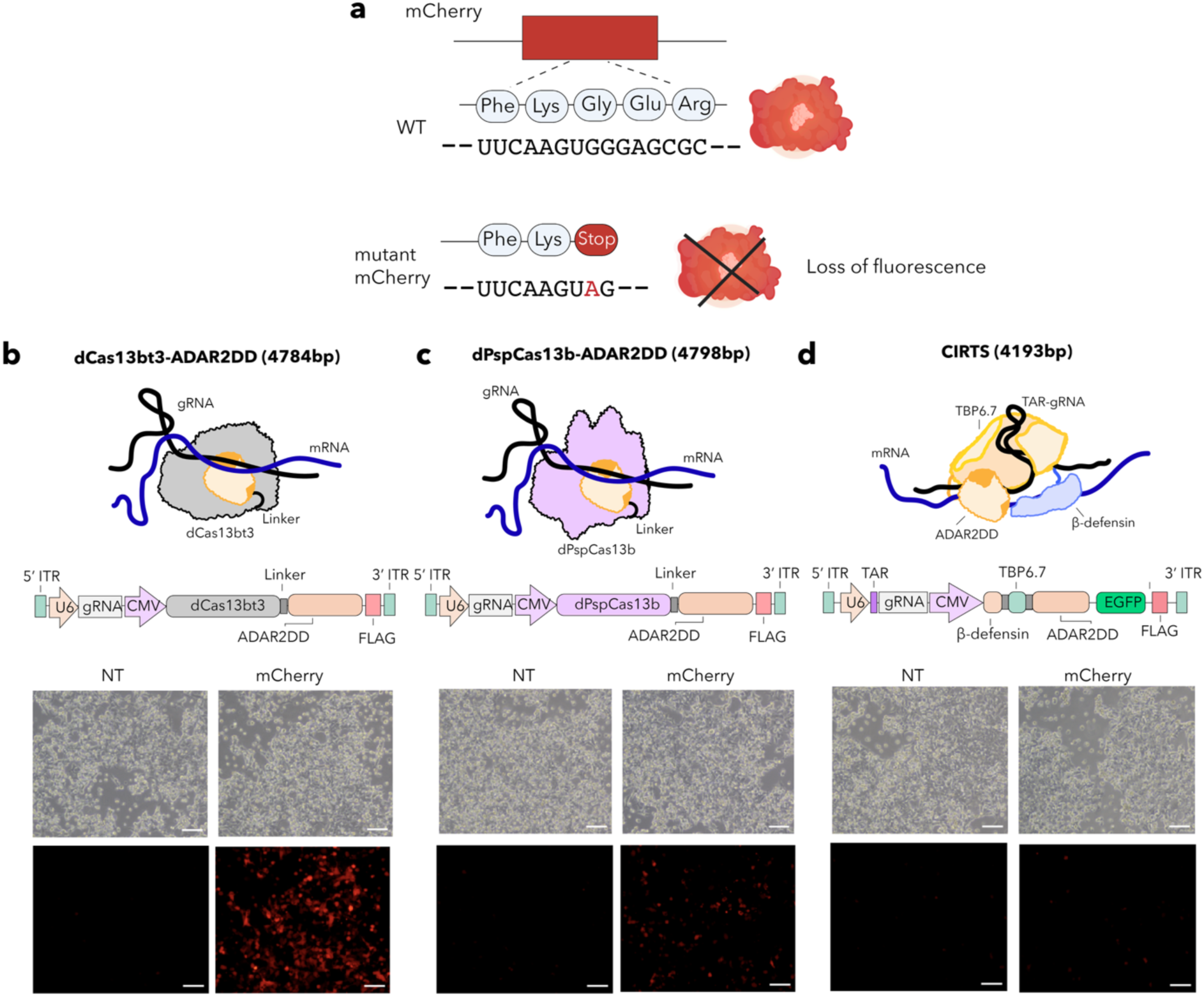
Comparison of all-in-one AAV RNA base editor designs with mCherry reporter assay. **(a)** Schematic of stop mutation in open reading frame of mCherry reporter leading to loss of fluorescence. Fluorescence images showing recovery of mCherry reporter from the **(b)** dCas13bt3-ADAR2DD, **(c)** dPspCas13b-ADAR2DD and **(d)** CIRTS base editors. dCas13bt3-ADAR2DD/NT: non-targeting gRNA, mCherry: dCas13bt3-ADAR2DD/mCherry targeting gRNA. Scale bar: 100 µm.

**Supplementary Figure 3.**
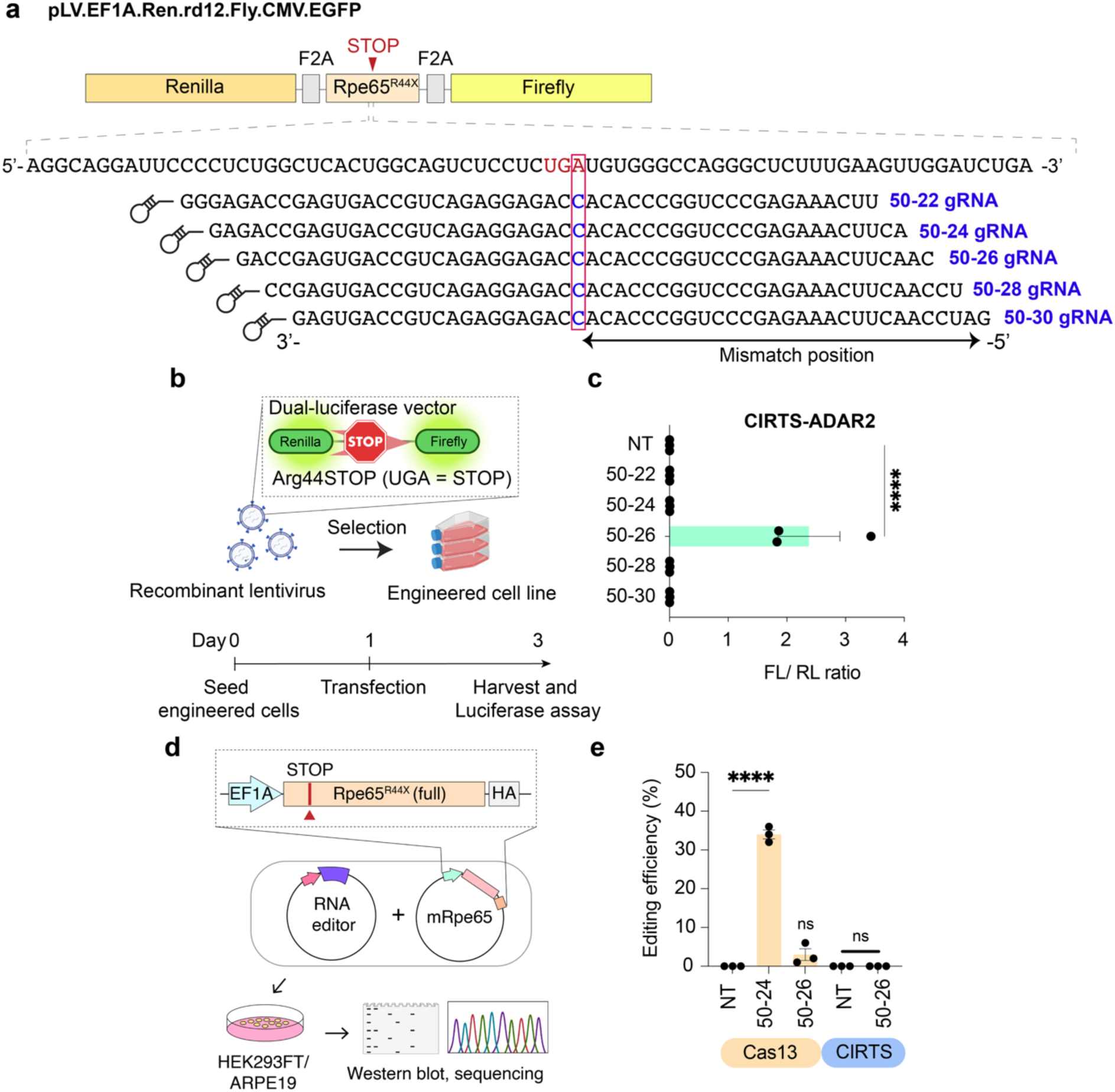
*In vitro* validation of compact RNA base editors. **(a)** Guide RNA screening with dual-luciferase assay. **(b)** Experimental procedure for dual-luciferase assay. **(c)** Recovery of firefly luciferase with CIRTS RNA base editor. FL: firefly luciferase, RL: renilla luciferase. **(d)** Experimental procedure for assessment of mRNA correction and protein recovery of RNA base editors. **(e)** Editing efficiency of dCas13bt3-ADAR2DD and CIRTS RNA base editors. Data presented as mean ± SEM. ****p < 0.0001, ns: not significant; One-Way ANOVA with Tukey’s multiple comparisons test (f and h). NT: non-targeting.

**Supplementary figure 4.**
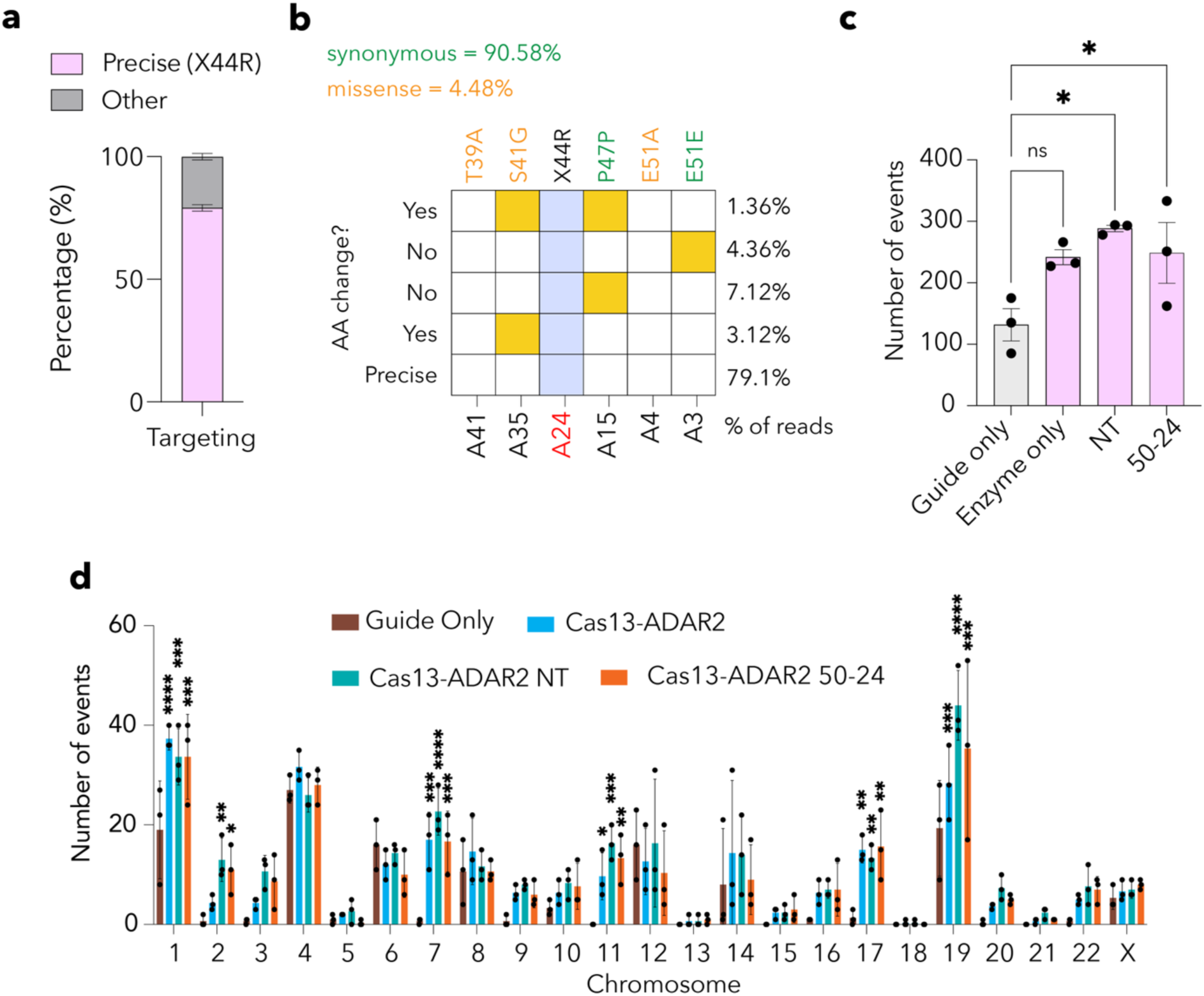
Bystander and off-target editing in ARPE19 cells. **(a)** Percentage of precise reads after base editing with dCas13bt3-ADAR2DD carrying the 50-24 gRNA in ARPE19 cells. **(b)** Heatmap showing proportion of reads with bystander edits that are either synonymous or non-synonymous. **(c)** Number of A-to-I editing events in ARPE19 cells across treatment groups and **(d)** individual chromosomes. Data presented as mean ± SEM. *p < 0.05, **p < 0.01, ***p < 0.001, ****p < 0.0001, ns: not significant; One-Way ANOVA with Tukey’s multiple comparisons test (c and d). Guide-only: 50-24 sgRNA only, Enzyme-only: dCas13bt3-ADAR2DD only, NT: dCas13bt3-ADAR2DD with non-targeting gRNA, 50-24: dCas13bt3-ADAR2DD with 50-24 gRNA.

**Supplementary figure 5.**
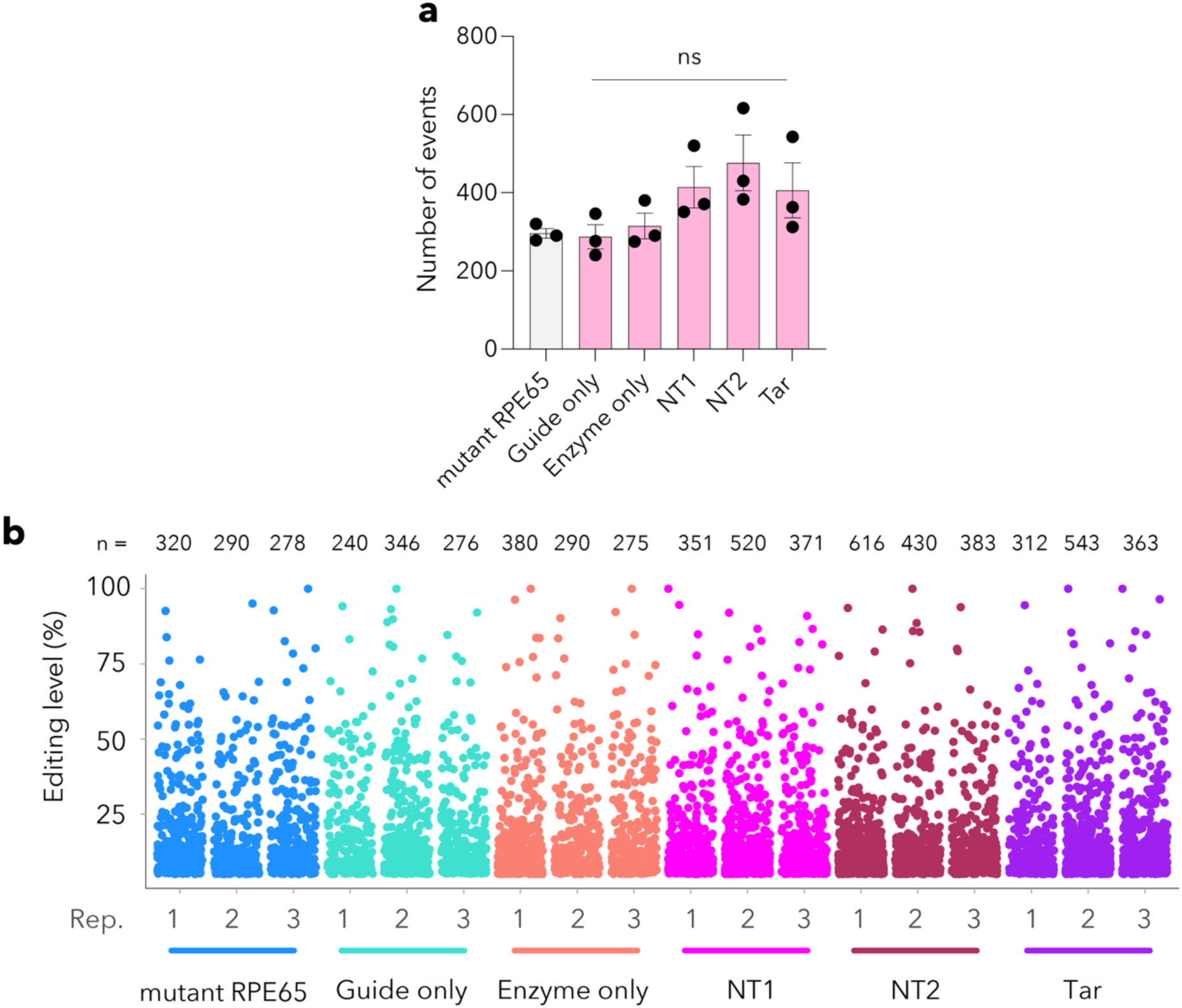
Comparison of different gRNAs with dCas13bt3-ADAR2DD RNA base editor in ARPE19 cells. **(a)** Barplot showing the total number of A-to-I events in each group. **(b)** Manhattan plot showing the number of A-to-I events in each biological replicate and the level of editing for each site. Data presented as mean ± SEM. ns: not significant; One-Way ANOVA with Tukey’s multiple comparisons test (a). NT: dCas13bt3-ADAR2DD with non-targeting gRNA, Tar: dCas13bt3-ADAR2DD with targeting gRNA, guide-only: targeting gRNA only, enzyme-only: dCas13bt3-ADAR2DD only.

**Supplementary figure 6.**
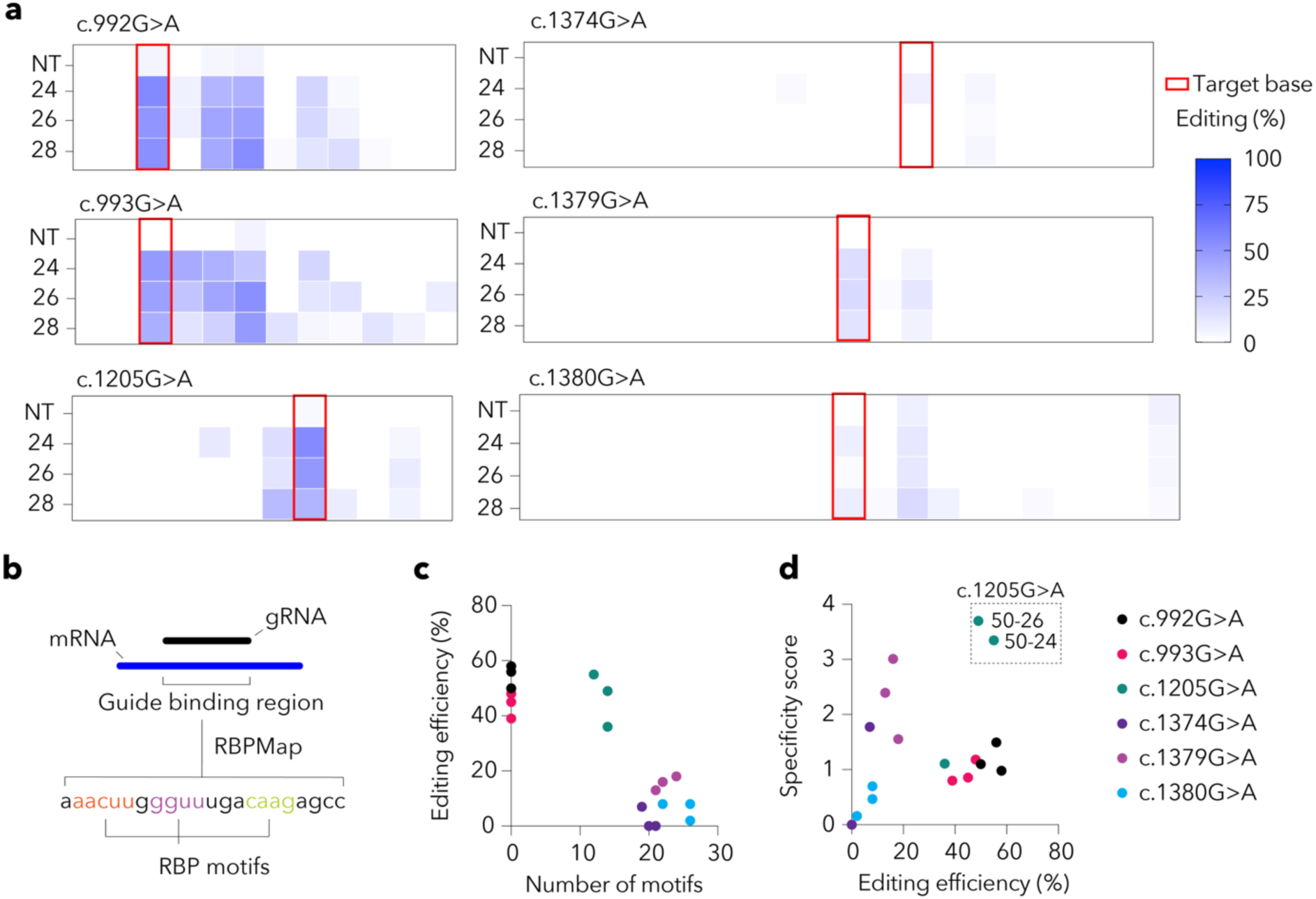
Bystander editing was observed with clinically reported RPE65 G>A mutations. **(a)** Heatmap showing the rate of bystander editing with each gRNA tested in the respective RPE65 G>A mutations. The red box indicates the target base. **(b)** Schematic of the procedure for identifying RNA binding protein (RBP) motifs in guide binding regions using RBPMap. **(c)** Editing efficiency was observed for each gRNA in relation to the respective mutations, as measured by the number of RBP motifs in that region. **(d)** Specificity scores for each gRNA in the respective mutations compared to their on-target efficiency. NT: non-targeting.

**Supplementary figure 7.**
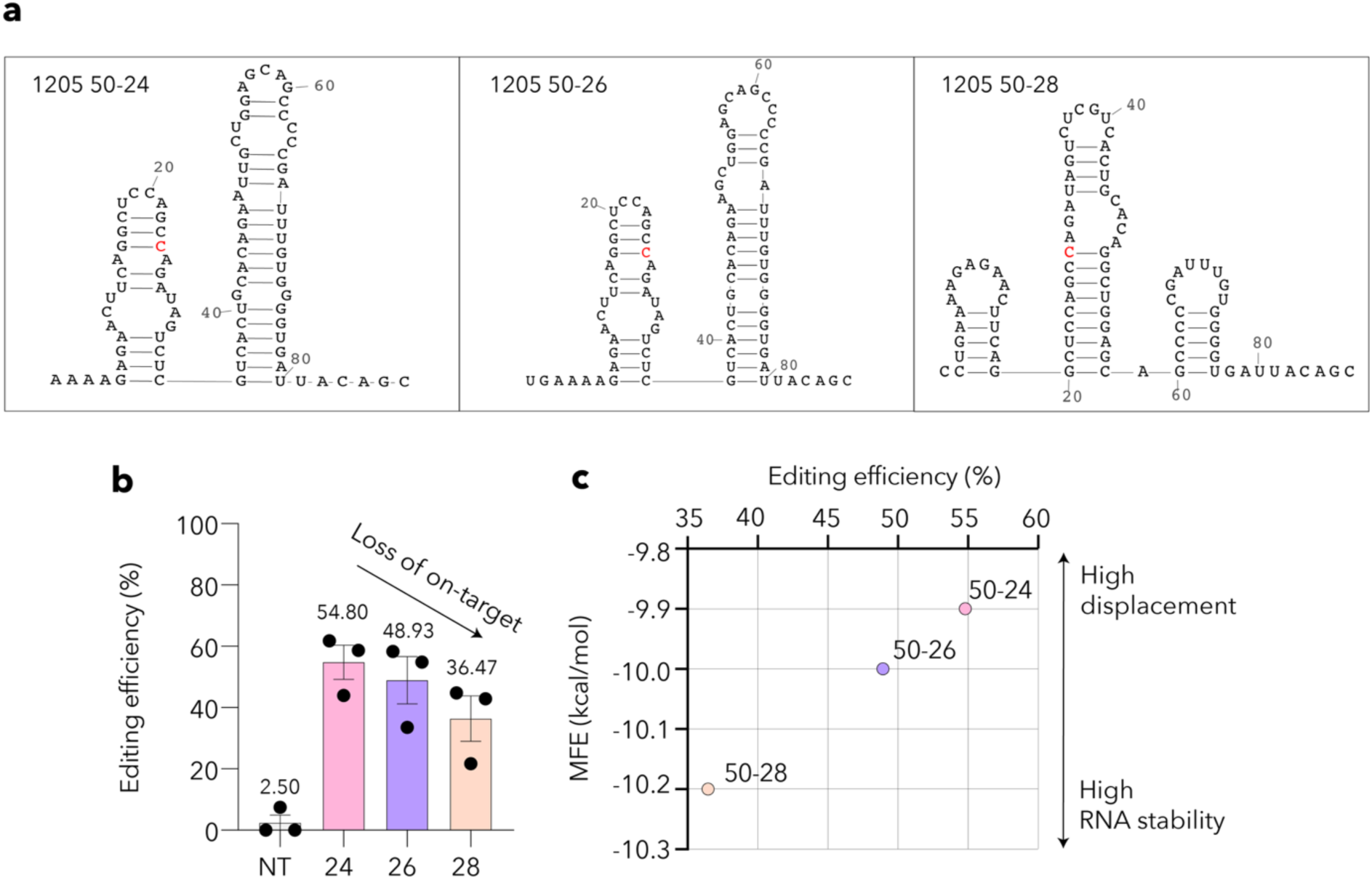
The impact of gRNA secondary structure on editing efficacy. **(a)** Secondary RNA structure of the c.1205G>A 50-24, 50-26 and 50-28 gRNAs with 3’ direct repeat. The 50-28 gRNA contains an additional stem loop at the 3’ end. Cytosine against target adenosine is highlighted in red. **(b)** Editing efficiency, determined from deep sequencing, shows a loss of on-target activity most prominently with the 50-28 gRNA. **(c)** The minimum free energy (MFE) of each gRNA secondary structure indicates the overall RNA stability achieved against on-target editing. Data presented as mean ± SEM. NT: non-targeting.

**Supplementary figure 8.**
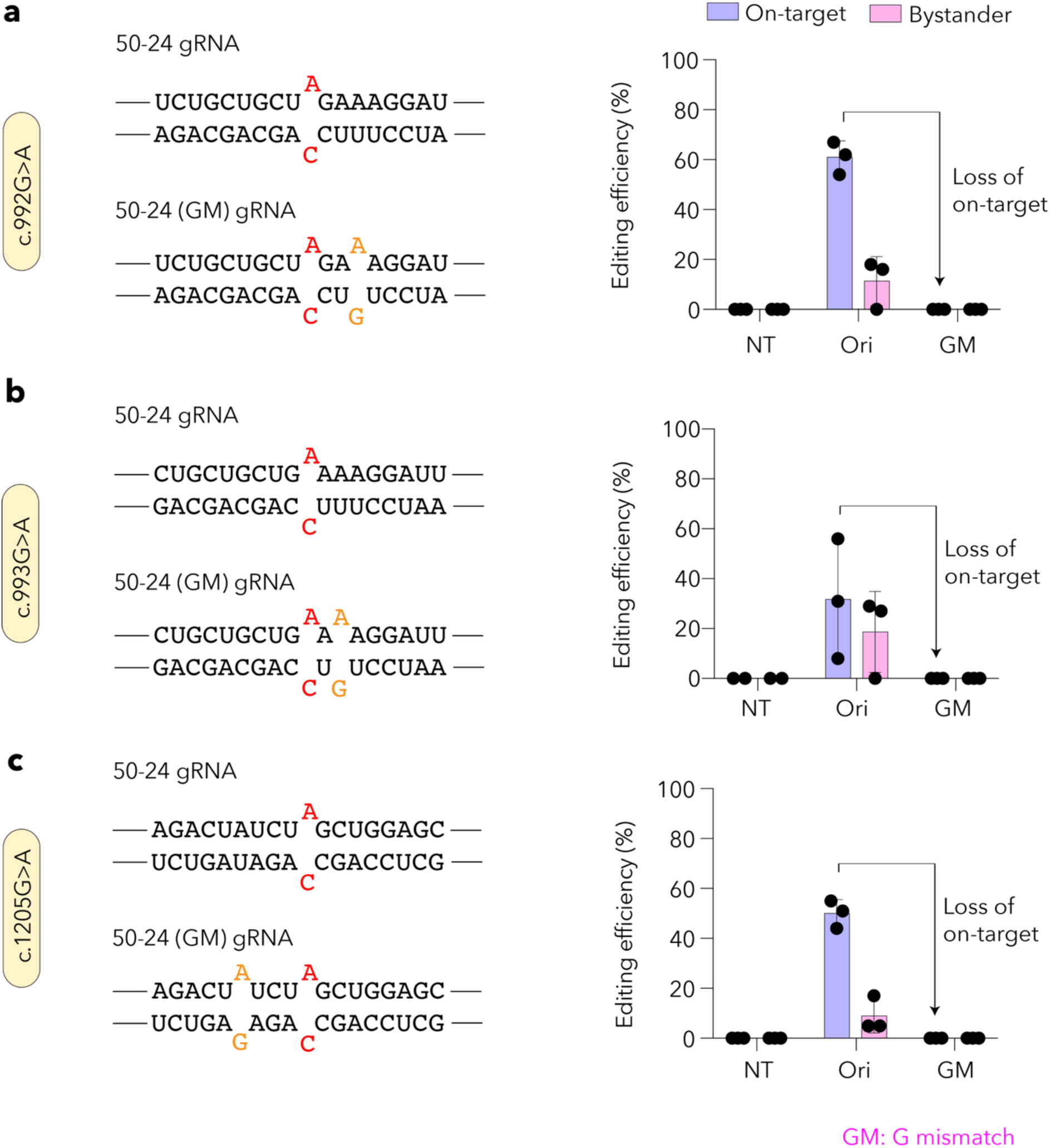
Validation of gRNA engineering with A:G mismatches against *RPE65* mutations. Schematic of original gRNA and gRNA with A:G mismatch (left) and editing efficiency observed with the respective gRNAs (right) for the **(a)** c.992G>A, **(b)** c.993G>A and **(c)** c.1205G>A mutations. Data presented as mean ± SEM. NT: non-targeting dRNA, Ori: original gRNA, GM: mismatch gRNA.

**Supplementary figure 9.**
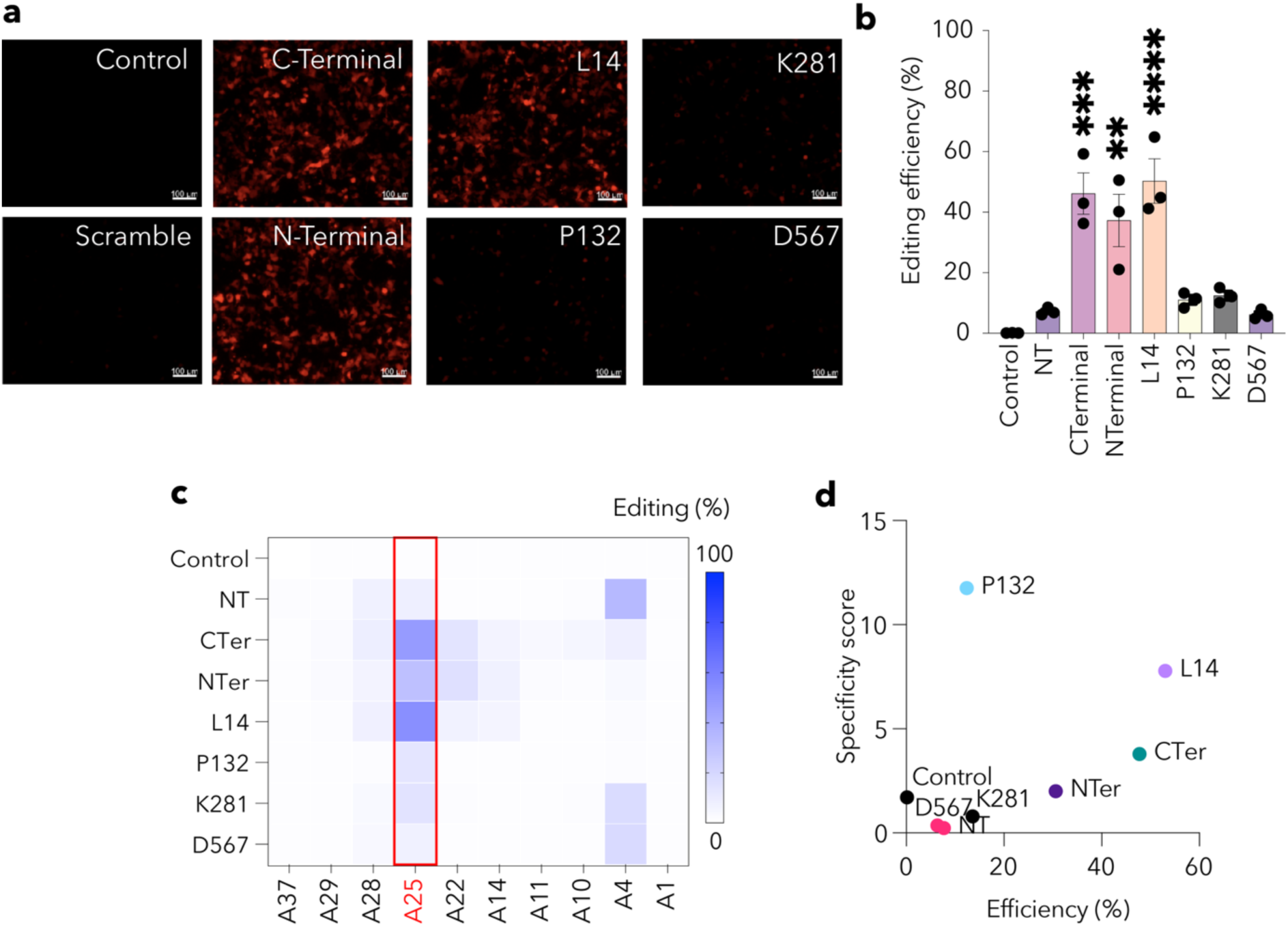
Screening of domain-inlaid dCas13bt3-ADAR2DD RNA base editors against mCherry reporter (with nonsense G>A mutant). **(a)** Fluorescence images showing recovery of mCherry reporter from domain-inlaid base editors. Scale bar: 100 μm **(b)** Editing efficiency determined from deep sequencing for the domain-inlaid base editors. **(c)** Heatmap for bystander editing observed for domain-inlaid base editors. **(d)** Specificity score for the domain-inlaid base editors against their on-target efficiency. Data presented as mean ± SEM. ****p < 0.0001, ns: not significant; One-Way ANOVA with Tukey’s multiple comparisons test (b). NT: non-targeting.

**Supplementary figure 10.**
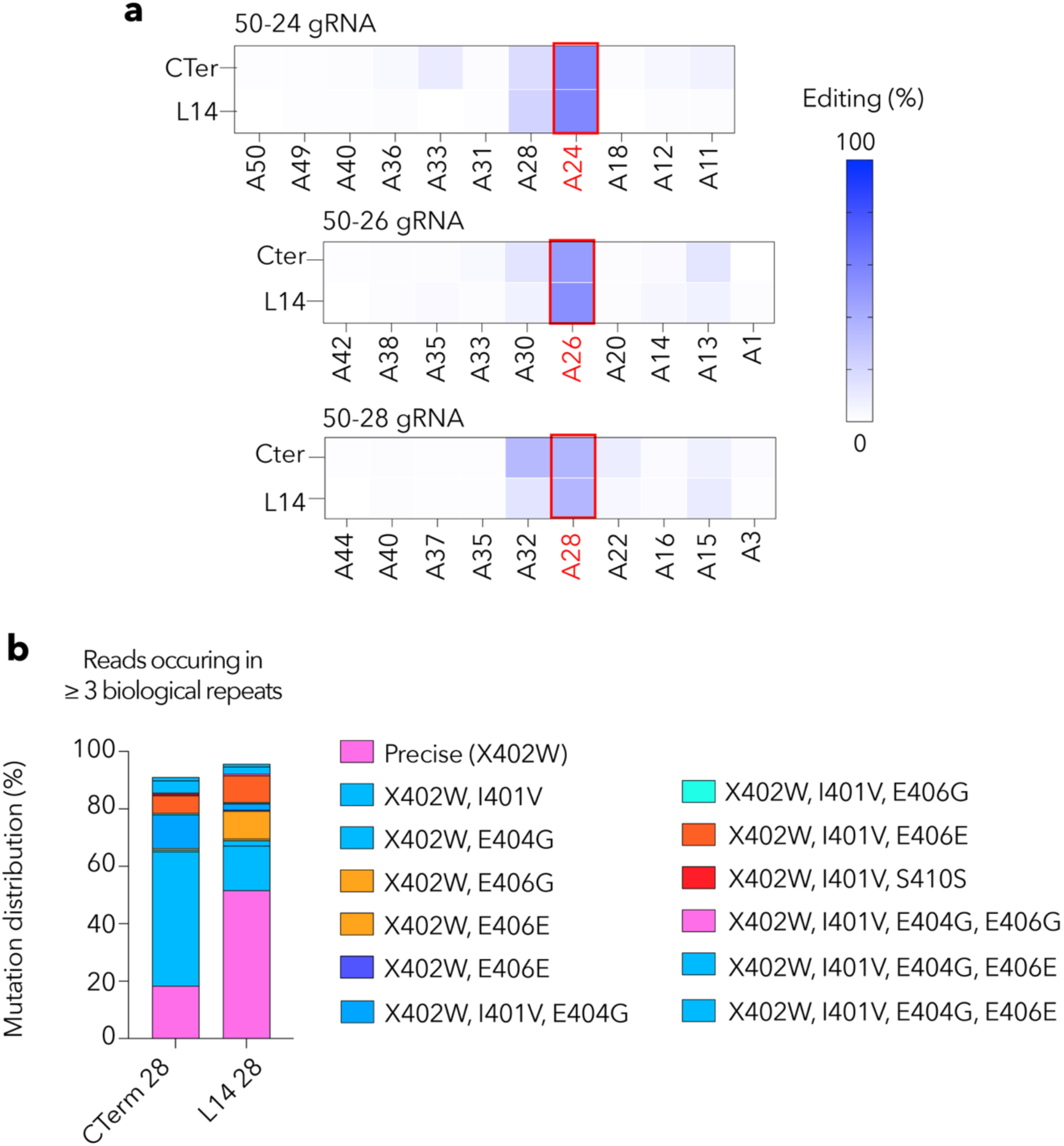
Bystander editing in *RPE65* c.1205G>A mutation with C-terminal and L14 base editors. **(a)** Heatmaps showing bystander editing profiles with panels of 50-24, 50-26, and 50-28 gRNAs between C-terminal and L14 base editors. **(b)** Mutation distribution of edited reads from C-terminal (C-Ter) and L14 base editors targeting the *RPE65* c.1205G>A mutation with 50-28 gRNAs.

**Supplementary figure 11.**
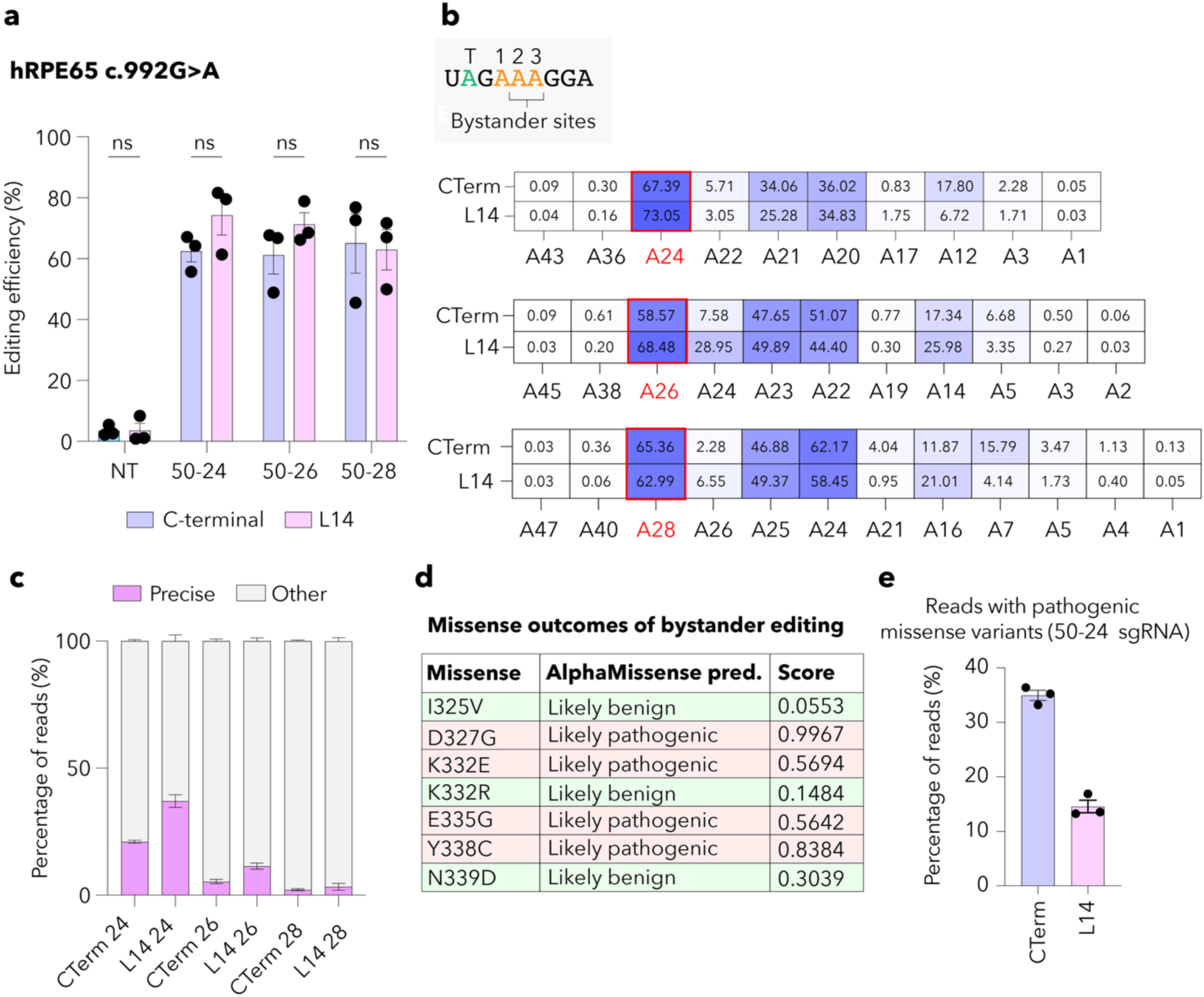
Validation of L14 base editor against *RPE65* c.992G>A mutation. **(a)** Editing efficiency observed with deep sequencing for the C-terminal (C-Term) and L14 base editor against the *RPE65* c.992G<A mutation. **(b)** Heatmap showing bystander editing with the C-terminal and L14 base editors with the different gRNAs tested. **(c)** Percentage of precise reads from the C-terminal and L14 base editors with the different gRNAs tested. **(d)** Table showing potential missense variants from bystander editing at the gRNA binding region. **(e)** Percentage of reads with likely pathogenic variants between C-terminal and L14 base editors with the 50-24 sgRNA. Data presented as mean ± SEM. ns: not significant; One-Way ANOVA with Tukey’s multiple comparisons test (a). NT: non-targeting.

**Supplementary figure 12.**
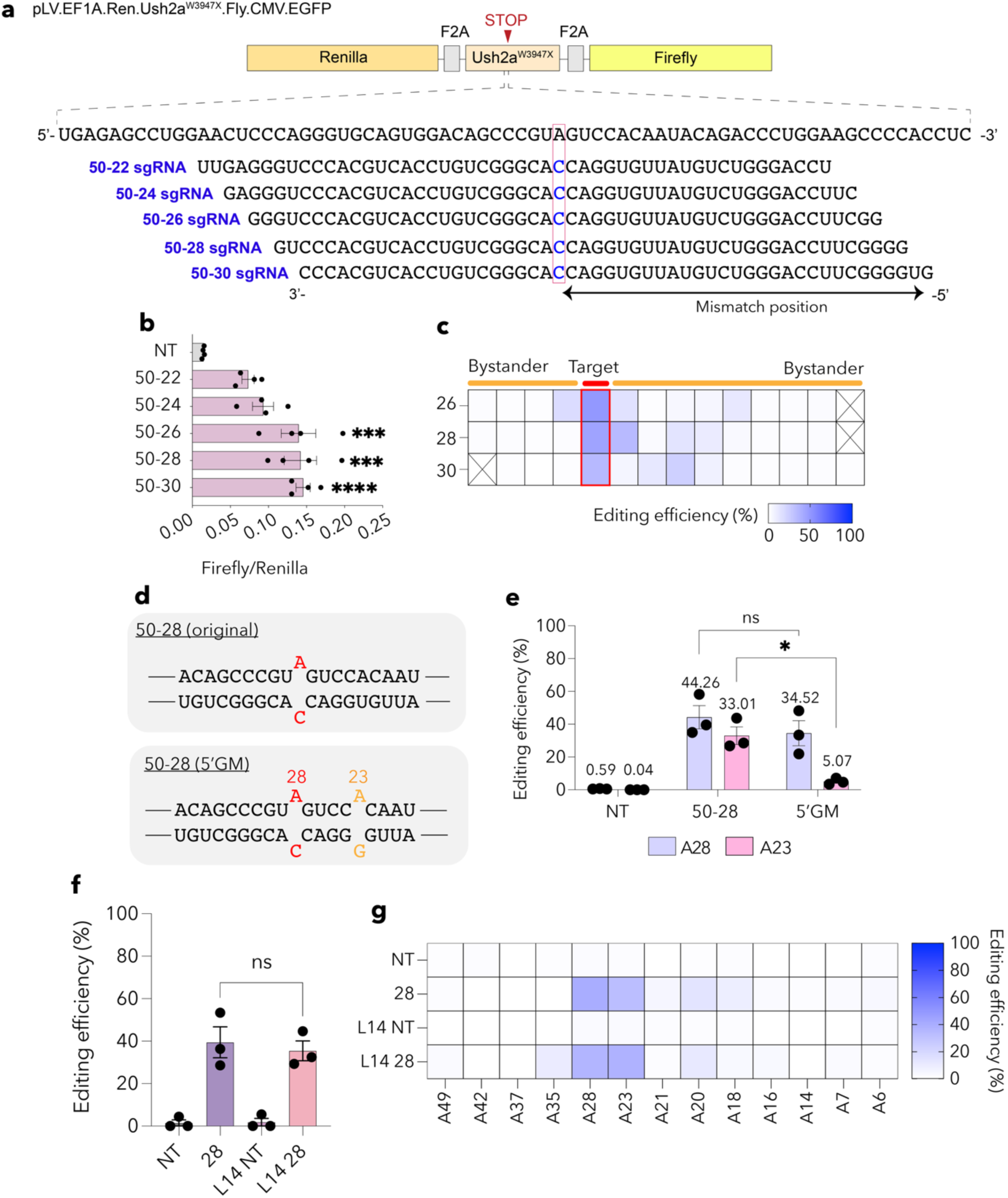
Validation of dCas13bt3-ADAR2DD base editor against *Ush2a*^W3947X^ mutation. **(a)** Schematic of gRNA screen against *Ush2a*^W3947X^ in reporter gene. The target base is indicated in the red dotted box. **(b)** Dual-luciferase assay showing the ratio of firefly to renilla luminescence, indicating the level of base correction at the target base with a panel of gRNAs (non-targeting [NT], 50-26, 50-28, and 50-30). **(c)** Heatmap showing the bystander profile at the *Ush2a*^W3947X^ target region. **(d)** Schematic showing original 50-28 gRNA and 50-28 gRNA with a G mismatch (GM) at A23. **(e)** Editing efficiency determined from deep sequencing of the original 50-28 and 50-28 (GM) gRNAs. **(f)** Editing efficiency determined from deep sequencing for C-terminal and L14 base editor carrying the 50-28 gRNA against *Ush2a*^W3947X^ mutation. **(g)** Heatmap showing bystander editing profile with C-terminal and L14 base editor with non-targeting and 50-28 gRNA. Data presented as mean ± SEM. *p < 0.05, ***p < 0.001, ****p < 0.0001, ns: not significant; One-Way ANOVA with Tukey’s multiple comparisons test (b, e and f).

**Supplementary figure 13.**
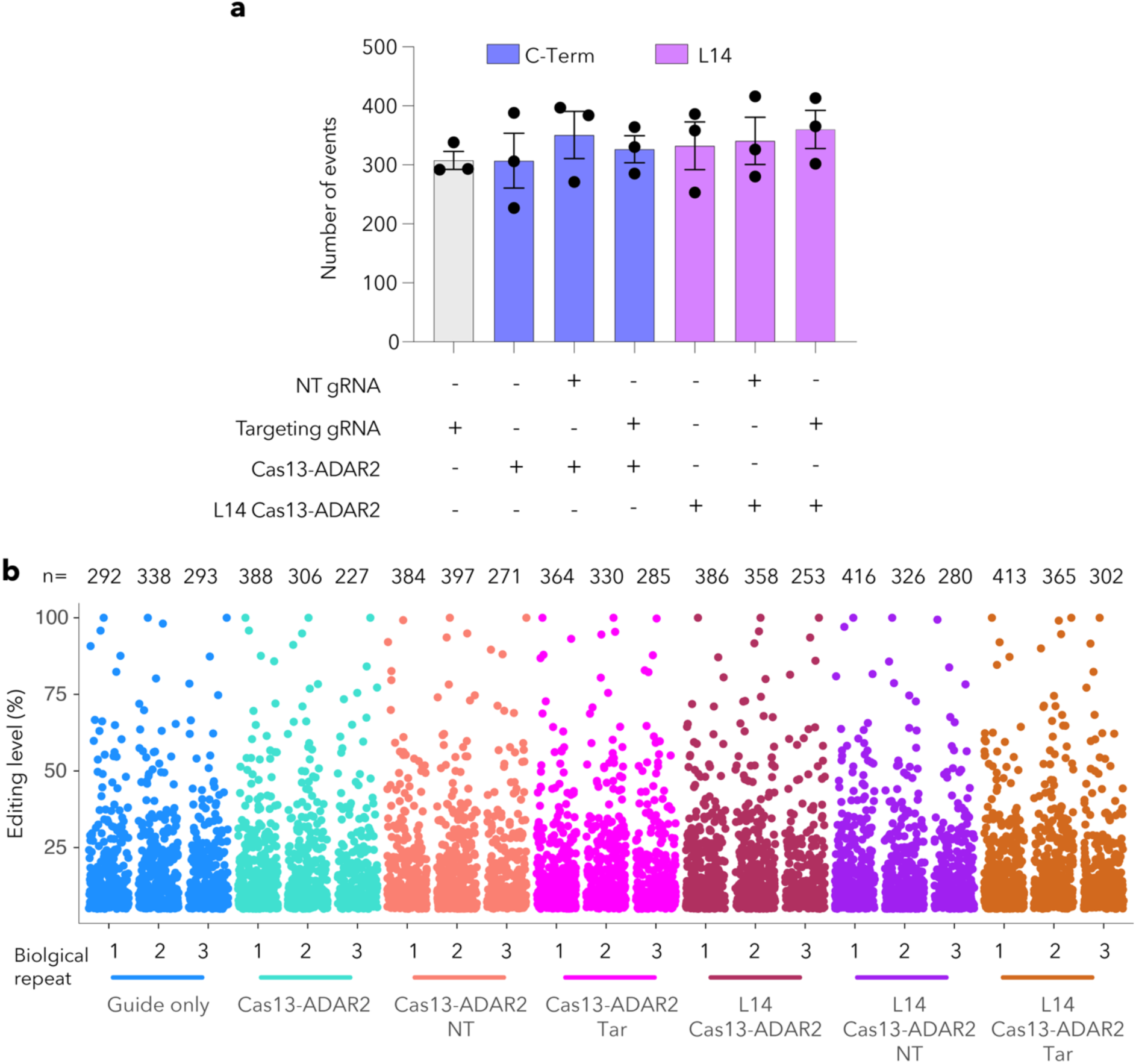
Transcriptome-wide A-to-I editing in HEK293FT cells from the C-terminal (C-Term) and L14 base editors. **(a)** Total number of A-to-I events in HEK293FT cells from gRNA (50-28 gRNA targeting *Ush2a*^W3947X^) only control, and C-terminal and L14 base editors with non-targeting and 50-28 gRNAs. **(b)** Manhattan plot showing A-to-I events in each biological replicate and rate of editing at each site. Data presented as mean ± SEM. NT: non-targeting gRNA, Tar: targeting gRNA.

**Supplementary figure 14.**
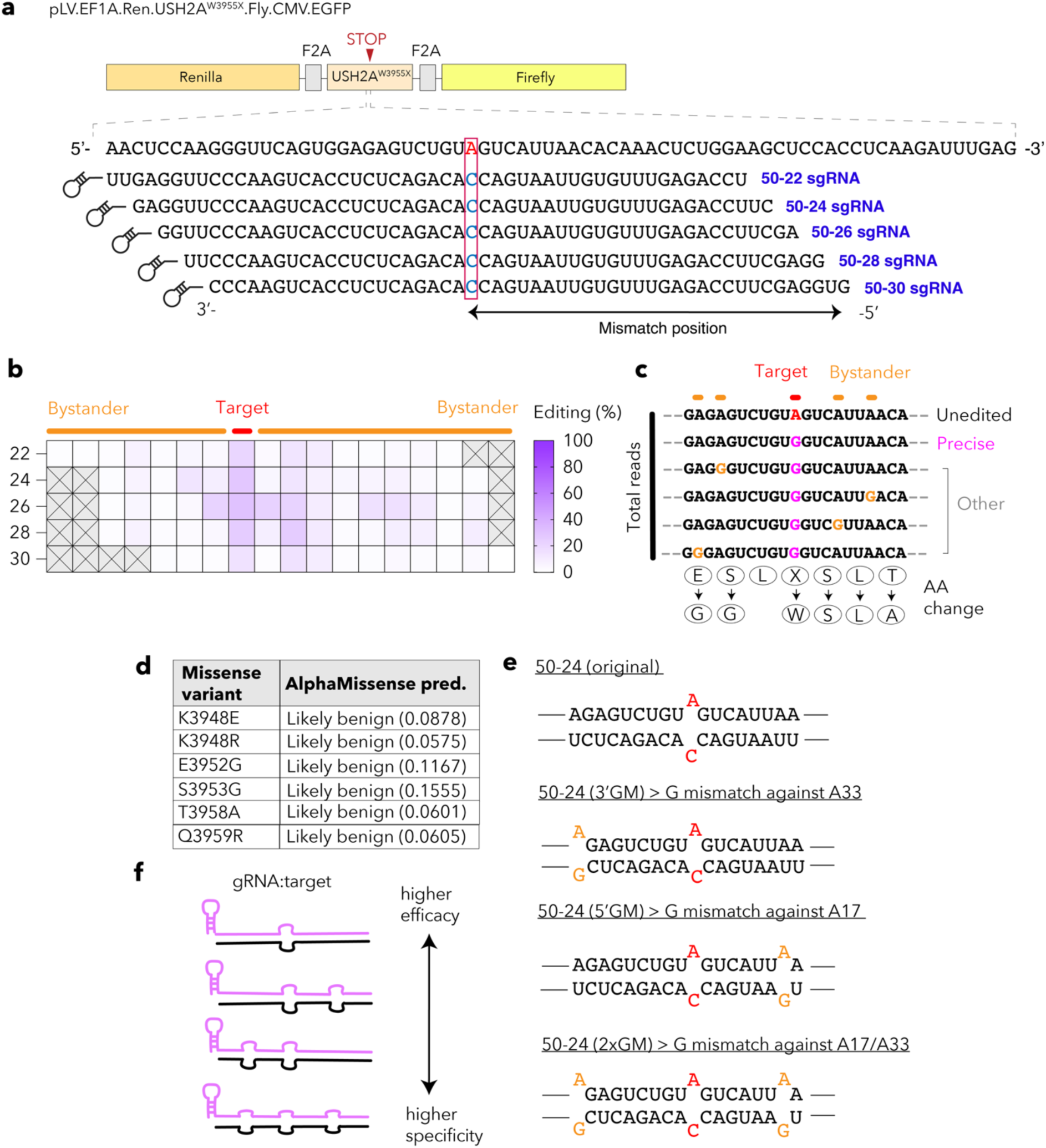
Validation of dCas13bt3-ADAR2DD base editor against the *USH2A*^W3955X^ mutation. **(a)** Schematic of *USH2A*^W3955X^ reporter gene along with a panel of gRNAs for screening. The target base is indicated in the red box. **(b)** Heatmap showing bystander editing profile with panel of gRNAs tested. **(c)** Schematic showing possible editing outcomes and impact on amino acid sequence from bystander editing. **(d)** Table showing potential missense variants in mRNA from base editing and their likelihood or pathogenicity from AlphaMissense, conservation and FoldX scores. **(e)** Schematic of mismatched gRNAs designed for reducing bystander editing showing original 50-24 gRNA, 50-24 (5’GM) gRNA, 50-24 (3’GM) gRNA, and 50-24 (2xGM) gRNA. **(f)** Schematic of rationale for incorporating mismatches into gRNA and potential impact on efficiency and specificity.

**Supplementary figure 15.**
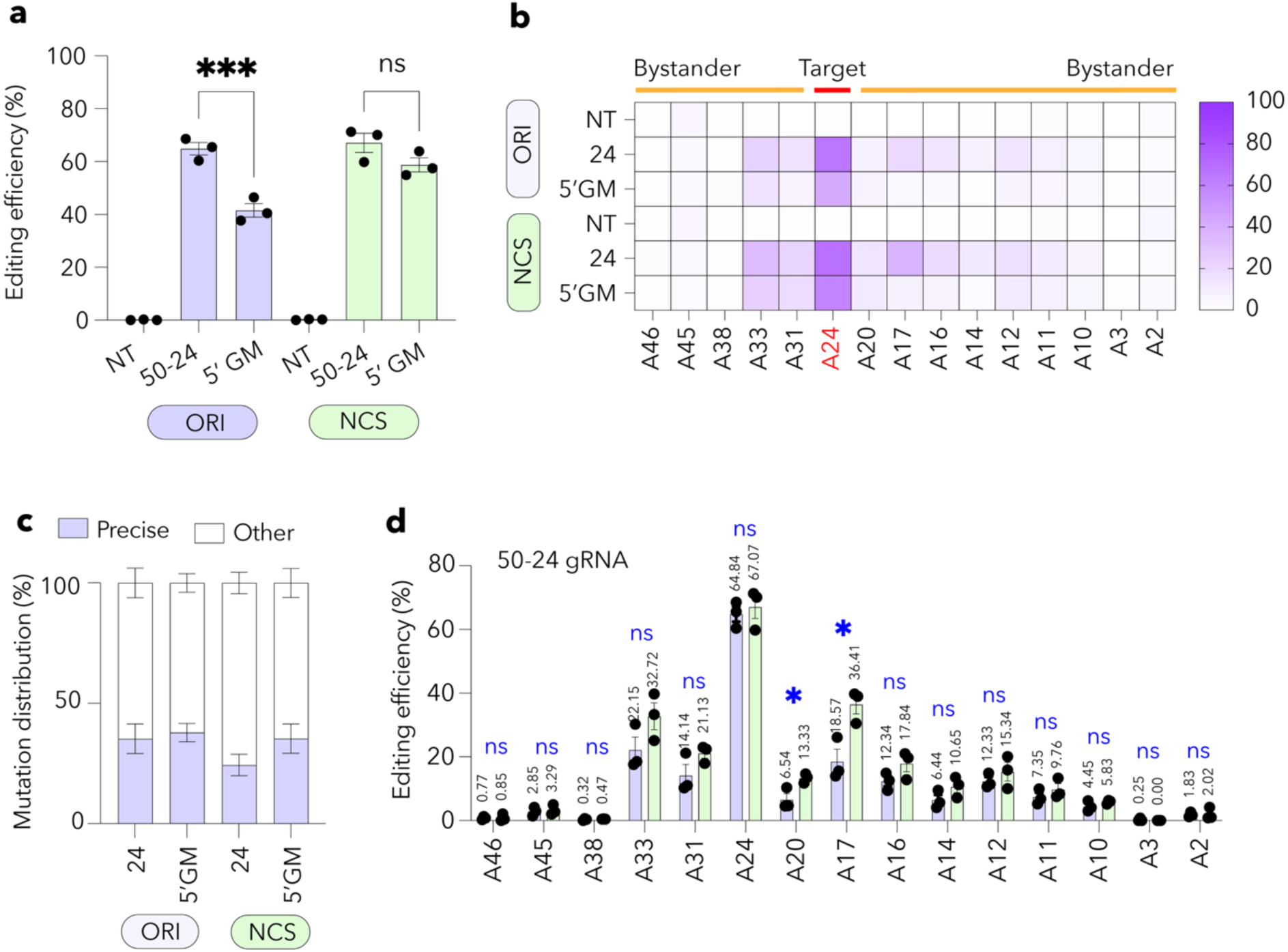
Efficacy and specificity of nucleocytoplasmic shuttling plasmids against *USH2A*^W3955X^ mutation. **(a)** Editing efficiency determined from deep sequencing for original and nucleocytoplasmic shuttling (NCS) constructs targeting the *USH2A*^W3955X^ mutation with non-targeting (NT), 50-24 and 50-24 (5’GM) gRNAs. **(b)** Heatmap showing bystander editing profile with original (ORI) and NCS constructs. **(c)** Percentage of precisely edited transcripts with original and NCS constructs. **(d)** Editing rates across gRNA binding region with 50-24 gRNA with original and NCS constructs. Data presented as mean ± SEM. *p < 0.05, ***p < 0.001, ns: not significant; One-Way ANOVA with Tukey’s multiple comparisons test (a and b).

**Supplementary figure 16.**
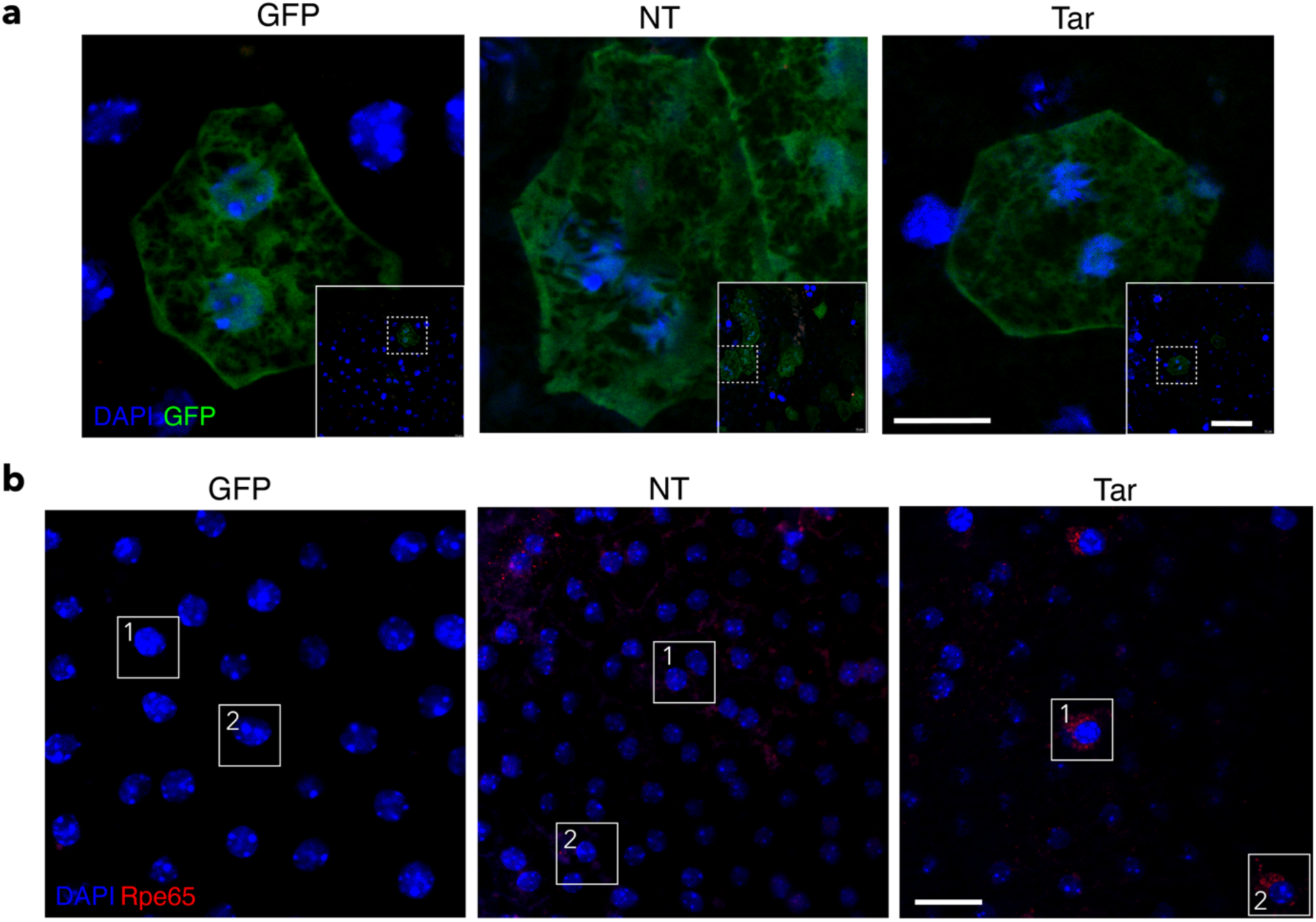
Immunofluorescence images from confocal microscopy of rd12 RPE flatmount. **(a)** Representative immunostaining showing GFP-expressing RPE cells in the rd12 RPE flatmount. Scale bar (left) 10 µm, (right) 50 µm. **(b)** Representative immunostaining showing expression of Rpe65 protein in the rd12 RPE flatmount. Scale bar: 20 µm NT: non-targeting AAV, Tar: *Rpe65*-targeting AAV.

**Supplementary figure 17.**
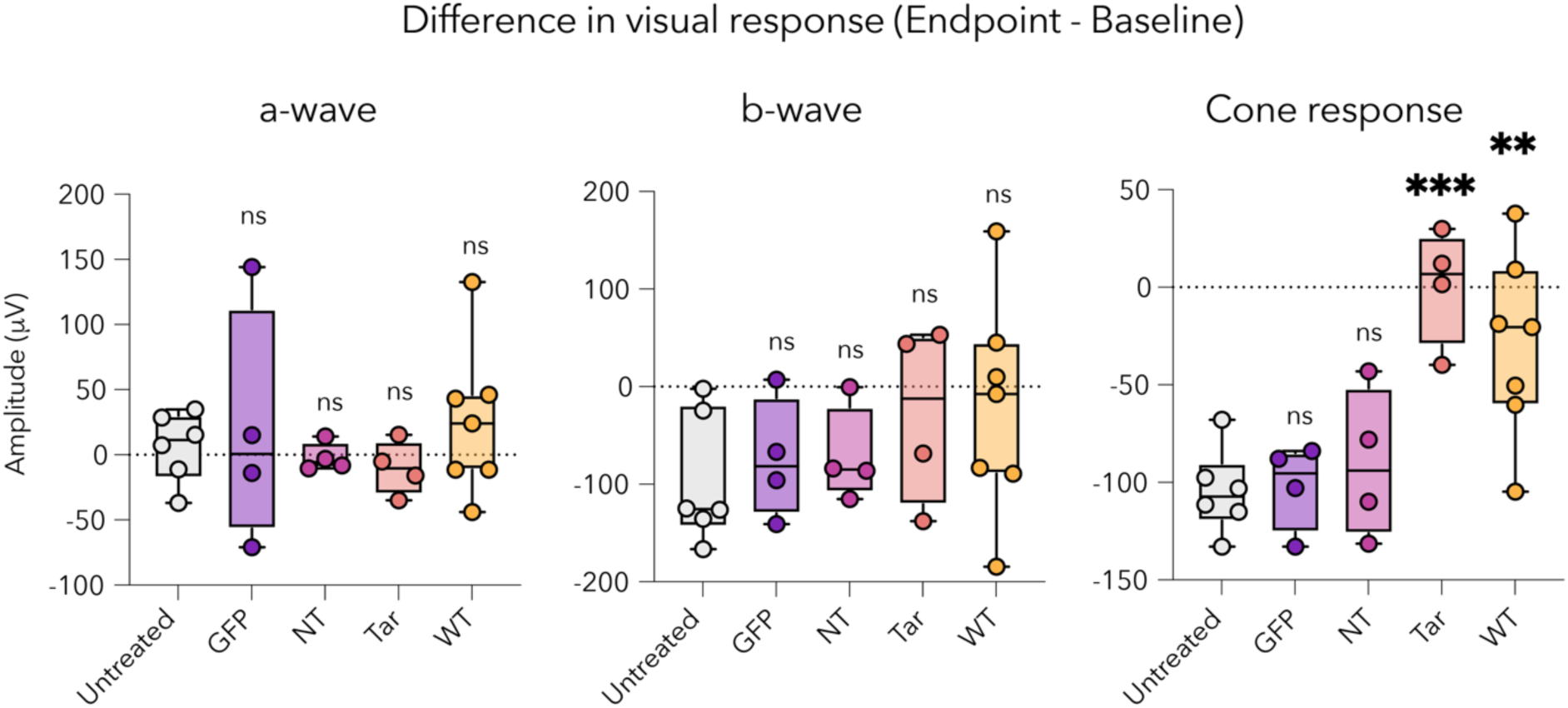
Difference in ERG measurements for a-wave, b-wave and cone responses in untreated rd12, rd12 treated with GFP, non-targeting (NT) and *Rpe65*-targeting (Tar) viruses, and wildtype mice. Data presented as mean ± SEM. **p < 0.01, ns: not significant; One-Way ANOVA with Tukey’s multiple comparisons test.

**Supplementary figure 18.**
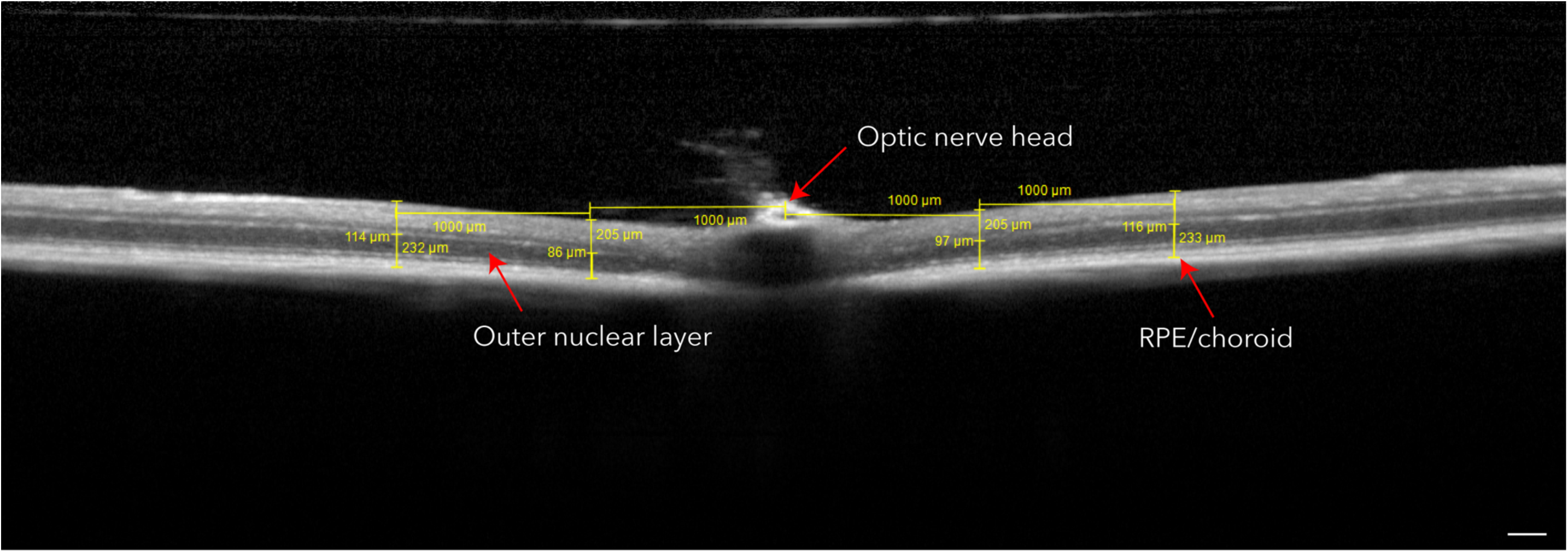
Representative OCT image from *Ush2a*^W3947X^ showing the measurements recorded to determine changes in retinal structure post AAV injections. Scale bar: 200 µm.

**Supplementary figure 19.**
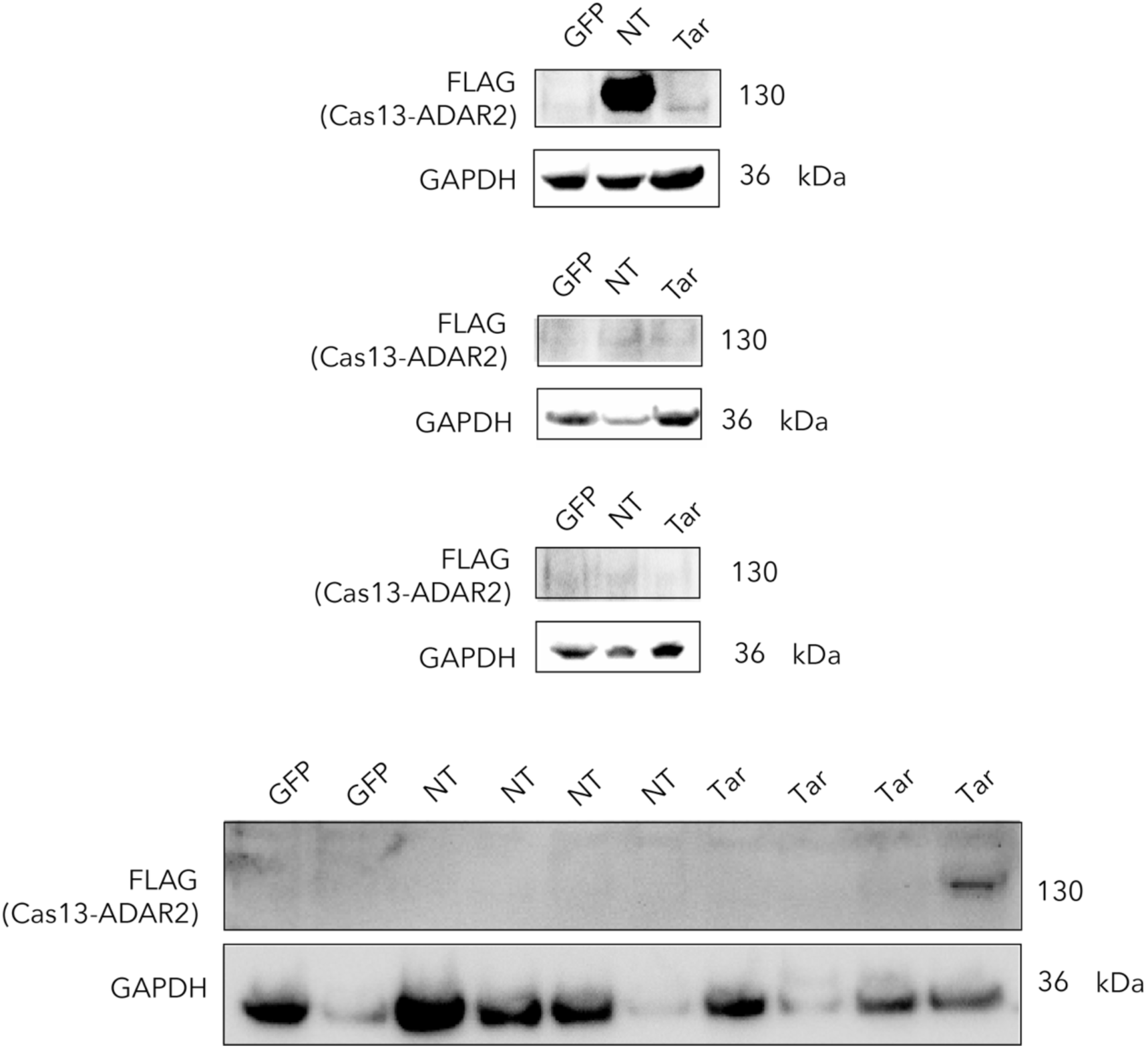
Western blot of *Ush2a*^W3947X^ mice retinae for FLAG expression, indicating delivery of dCas13bt3-ADAR2DD into retina. NT: non-targeting gRNA, Tar: *Ush2a*-targeting gRNA.

**Supplementary figure 20.**
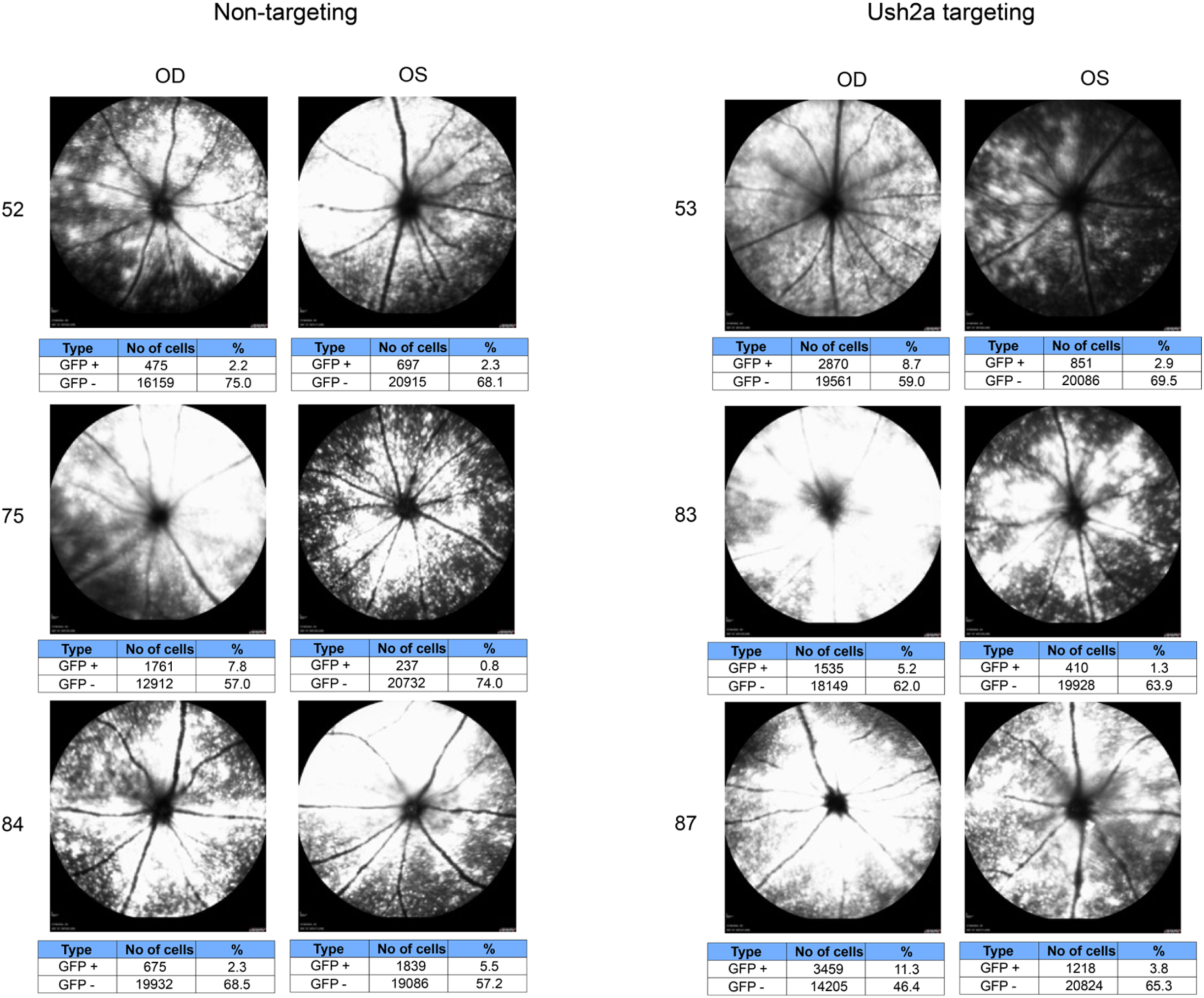
Fluorescence activated cell sorting (FACS) based on GFP expression from Ush2a retinae post dissociation.

**Supplementary figure 21.**
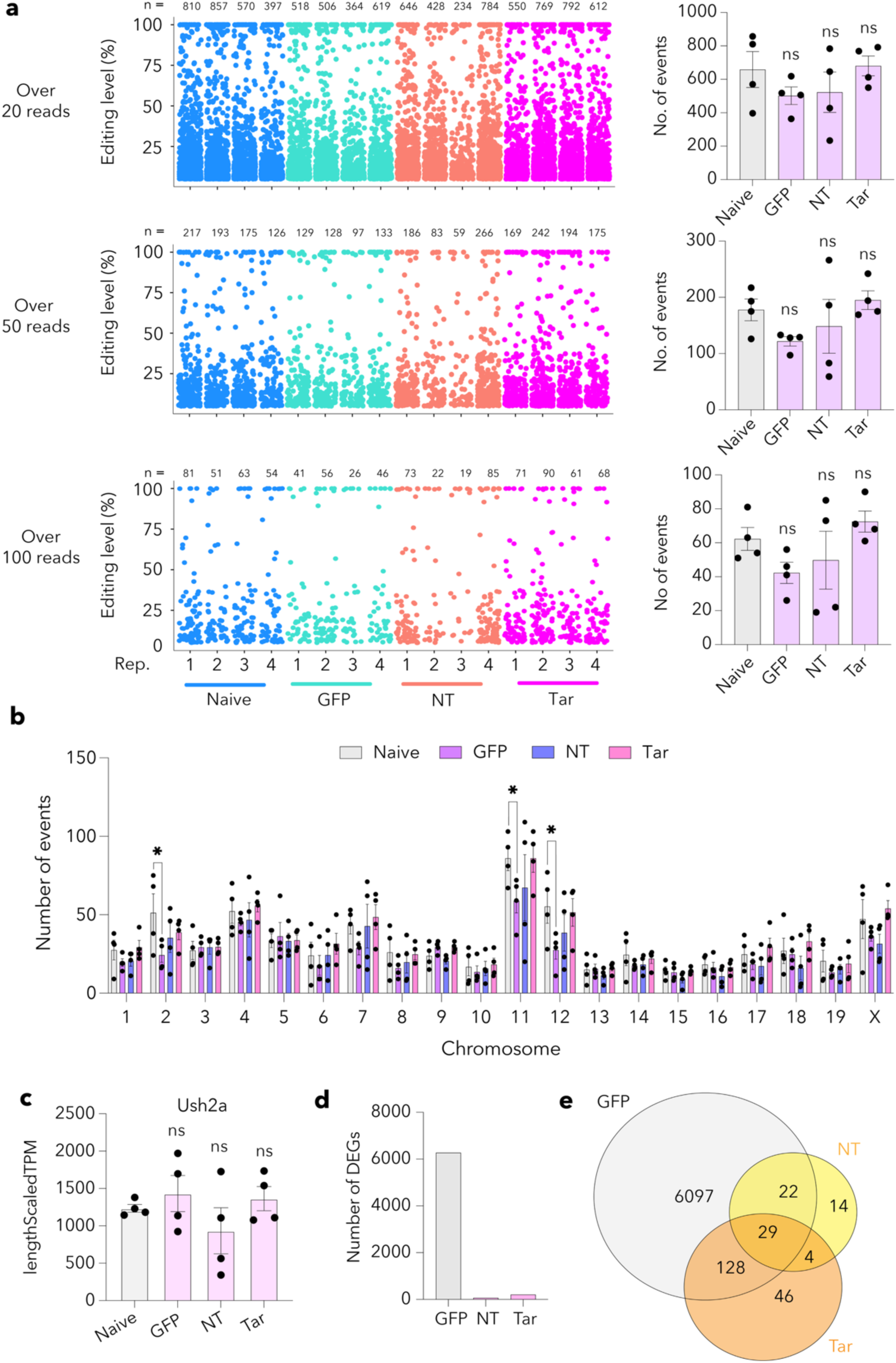
RNA sequencing of *Ush2a*^W3947X^ mice retinae following treatment with dCas13bt3-ADAR2DD. **(a)** Manhattan and bar plots showing the total number of A-to-I editing events across naïve (untreated), GFP-only, non-targeting (NT), and *Ush2a*-targeting (Tar) groups. **(b)** A-to-I events across the mouse chromosomes. **(c)** Transcript levels of *Ush2a* RNA from the different treatment groups. **(d)** Total level of DEGs compared to naïve mice. **(e)** Venn diagram showing the intersection of DEGs across the treatment groups. Data presented as mean ± SEM. *p < 0.05, ns: not significant; One-Way ANOVA with Tukey’s multiple comparisons test (a, b and c).

## REFERENCES

1. Botto C, Rucli M, Tekinsoy MD, Pulman J, Sahel J-A, Dalkara D. Early and late stage gene therapy interventions for inherited retinal degenerations. Prog Retin Eye Res 86, 100975 (2022).

2. Berger W, Kloeckener-Gruissem B, Neidhardt J. The molecular basis of human retinal and vitreoretinal diseases. Prog Retin Eye Res 29, 335–375 (2010).

3. Fry LE, Mcclements ME, Maclaren RE. Analysis of Pathogenic Variants Correctable With CRISPR Base Editing Among Patients With Recessive Inherited Retinal Degeneration. JAMA Ophthalmol, 139 (2021).

4. Fontana M, et al. CRISPR-Cas9 Gene Editing with Nexiguran Ziclumeran for ATTR Cardiomyopathy. N Engl J Med 391, 2231–2241 (2024).

5. Musunuru K, et al. In vivo CRISPR base editing of PCSK9 durably lowers cholesterol in primates. Nature 593, 429–434 (2021).

6. Amoasii L, et al. Gene editing restores dystrophin expression in a canine model of Duchenne muscular dystrophy. Science 362, 86–91 (2018).

7. Jang H, et al. Application of prime editing to the correction of mutations and phenotypes in adult mice with liver and eye diseases. Nature Biomedical Engineering 6, 181–194 (2022).

8. Gaudelli NM, et al. Programmable base editing of A•T to G•C in genomic DNA without DNA cleavage. Nature 551, 464–471 (2017).

9. Komor AC, Kim YB, Packer MS, Zuris JA, Liu DR. Programmable editing of a target base in genomic DNA without double-stranded DNA cleavage. Nature 533, 420–424 (2016).

10. Anzalone AV, et al. Search-and-replace genome editing without double-strand breaks or donor DNA. Nature 576, 149–157 (2019).

11. Xiao Q, et al. Rescue of autosomal dominant hearing loss by in vivo delivery of mini dCas13X-derived RNA base editor. Sci Transl Med 14, eabn0449 (2022).

12. Xue Y, et al. RNA base editing therapy cures hearing loss induced by *OTOF* gene mutation. Mol Ther 31, 3520–3530 (2023).

13. Abudayyeh OO, et al. C2c2 is a single-component programmable RNA-guided RNA-targeting CRISPR effector. Science 353, aaf5573 (2016).

14. Kannan S, et al. Compact RNA editors with small Cas13 proteins. Nature Biotechnology, 1-4 (2021).

15. Xu C, et al. Programmable RNA editing with compact CRISPR–Cas13 systems from uncultivated microbes. Nat Methods 18, 499–506 (2021).

16. Cox DBT, et al. RNA editing with CRISPR-Cas13. Science 358, 1019 (2017).

17. Rauch S, He E, Srienc M, Zhou H, Zhang Z, Dickinson BC. Programmable RNA-Guided RNA Effector Proteins Built from Human Parts. Cell 178, 122–134.e112 (2019).

18. Pang JJ, et al. Retinal degeneration 12 (rd12): a new, spontaneously arising mouse model for human Leber congenital amaurosis (LCA). Mol Vis 11, 152–162 (2005).

19. Fu Z-C, Gao B-Q, Nan F, Ma X-K, Yang L. DEMINING: A deep learning model embedded framework to distinguish RNA editing from DNA mutations in RNA sequencing data. Genome Biol 25, 258 (2024).

20. Wang X, et al. Develop a Compact RNA Base Editor by Fusing ADAR with Engineered EcCas6e. Advanced Science 10, 2206813 (2023).

21. Behroozi J, Shahbazi S, Bakhtiarizadeh MR, Mahmoodzadeh H. Genome-Wide Characterization of RNA Editing Sites in Primary Gastric Adenocarcinoma through RNA-seq Data Analysis. International Journal of Genomics 2020, 6493963 (2020).

22. Eggington JM, Greene T, Bass BL. Predicting sites of ADAR editing in double-stranded RNA. Nature Communications 2, 319 (2011).

23. Corley M, Burns MC, Yeo GW. How RNA-Binding Proteins Interact with RNA: Molecules and Mechanisms. Mol Cell 78, 9–29 (2020).

24. Wang Y, et al. A circularly permuted CasRx platform for efficient, site-specific RNA editing. Nature Biotechnology, (2024).

25. Ofer K, Benjamin BL, Owen RSD, Aartjan JWtV, Cameron M. RNA structure modulates Cas13 activity and enables mismatch detection. bioRxiv, 2023.2010.2005.560533 (2023).

26. Liu Y, et al. REPAIRx, a specific yet highly efficient programmable A > I RNA base editor. The EMBO Journal 39, e104748 (2020).

27. Nguyen Tran MT, et al. Engineering domain-inlaid SaCas9 adenine base editors with reduced RNA off-targets and increased on-target DNA editing. Nature Communications 11, 4871 (2020).

28. Cheng J, et al. Accurate proteome-wide missense variant effect prediction with AlphaMissense. Science 381, eadg7492 (2023).

29. Fry LE, et al. Comparison of CRISPR-Cas13b RNA base editing approaches for USH2A-associated inherited retinal degeneration. Commun Biol 8, 200 (2025).

30. Gruber C, et al. Engineered, nucleocytoplasmic shuttling Cas13d enables highly efficient cytosolic RNA targeting. Cell Discovery 10, 42 (2024).

31. Seeliger MW, et al. New views on RPE65 deficiency: the rod system is the source of vision in a mouse model of Leber congenital amaurosis. Nat Genet 29, 70–74 (2001).

32. Elenius V. Cone and Rod Activity in the Electroretinogram Evoked by Double Flashes of Light. Archives of Ophthalmology 81, 618–621 (1969).

33. Katrekar D, et al. In vivo RNA editing of point mutations via RNA-guided adenosine deaminases. Nat Methods 16, 239–242 (2019).

34. Katrekar D, et al. Efficient in vitro and in vivo RNA editing via recruitment of endogenous ADARs using circular guide RNAs. Nature Biotechnology, (2022).

35. Li G, et al. Mini-dCas13X–mediated RNA editing restores dystrophin expression in a humanized mouse model of Duchenne muscular dystrophy. The Journal of Clinical Investigation 133, (2023).

36. Yi Z, et al. Engineered circular ADAR-recruiting RNAs increase the efficiency and fidelity of RNA editing in vitro and in vivo. Nature Biotechnology 40, 946–955 (2022).

37. Hołubowicz R, et al. Safer and efficient base editing and prime editing via ribonucleoproteins delivered through optimized lipid-nanoparticle formulations. Nature Biomedical Engineering 9, 57–78 (2025).

38. Choi EH, et al. In vivo base editing rescues cone photoreceptors in a mouse model of early-onset inherited retinal degeneration. Nature Communications 13, 1830 (2022).

39. Muller A, et al. High-efficiency base editing in the retina in primates and human tissues. Nat Med 31, 490–501 (2025).

40. Bennett J, et al. AAV2 Gene Therapy Readministration in Three Adults with Congenital Blindness. Sci Transl Med 4, 120ra115 (2012).

41. Moreno AM, et al. Immune-orthogonal orthologues of AAV capsids and of Cas9 circumvent the immune response to the administration of gene therapy. Nat Biomed Eng 3, 806–816 (2019).

42. Sun Y, et al. Improved RNA base editing with guide RNAs mimicking highly edited endogenous ADAR substrates. Nature Biotechnology, (2025).

43. Li H, et al. Engineering a photoactivatable A-to-I RNA base editor for gene therapy in vivo. Nature Biotechnology, (2025).

44. Li G, et al. Specific and efficient RNA A-to-I editing through cleavage of an ADAR inhibitor. Nature Biotechnology, (2025).

45. Stephenson JD, Totoo P, Burke David F, Jänes J, Beltrao P, Martin Maria J. ProtVar: mapping and contextualizing human missense variation. Nucleic Acids Research 52, W140–W147 (2024).

46. Lorenz R, et al. ViennaRNA Package 2.0. Algorithms for Molecular Biology 6, 26 (2011).

47. Patro R, Duggal G, Love MI, Irizarry RA, Kingsford C. Salmon provides fast and bias-aware quantification of transcript expression. Nat Methods 14, 417–419 (2017).

48. Love MI, Huber W, Anders S. Moderated estimation of fold change and dispersion for RNA-seq data with DESeq2. Genome Biol 15, 550 (2014).

49. Dureau P, et al. Quantitative analysis of subretinal injections in the rat. Graefe’s Archive for Clinical and Experimental Ophthalmology 238, 608–614 (2000).

